# Cryo-EM structures reveal ubiquinone-10 binding to mitochondrial complex I and conformational transitions associated with Q-site occupancy

**DOI:** 10.1101/2022.02.11.480065

**Authors:** Injae Chung, John J. Wright, Hannah R. Bridges, Bozhidar S. Ivanov, Olivier Biner, Caroline S. Pereira, Guilherme M. Arantes, Judy Hirst

**Affiliations:** MRC Mitochondrial Biology Unit, University of Cambridge, The Keith Peters Building, Cambridge Biomedical Campus, Hills Road, Cambridge, CB2 0XY, UK; Department of Biochemistry, Instituto de Química, Universidade de São Paulo, Av. Prof. Lineu Prestes 748, 05508-900, São Paulo, SP, Brazil; Institute of Plant and Microbial Biology, University of Zurich, Zollikerstrasse 107, 8008 Zürich

## Abstract

Mitochondrial complex I is a central metabolic enzyme that uses the reducing potential of NADH to reduce ubiquinone-10 (Q10) and drive four protons across the inner mitochondrial membrane, powering oxidative phosphorylation. Although many complex I structures are now available, structures of Q10-bound states have remained elusive. Here, we reconstitute mammalian complex I into phospholipid nanodiscs with exogenous Q10. Using cryo-EM, we reveal a Q10 molecule occupying the full length of the Q-binding site in the ‘active’ (ready-to-go) resting state (plus a matching substrate-free structure) and apply molecular dynamics simulations to propose how the charge states of key residues influence the Q10 binding pose. By comparing ligand-bound and ligand-free forms of the ‘deactive’ resting state (that require reactivating to catalyse), we begin to define how substrate binding restructures the deactive Q-binding site, providing insights into its physiological and mechanistic relevance.

## Introduction

Mitochondrial complex I (NADH:ubiquinone oxidoreductase) is an intricate ∼1 MDa multimeric membrane-bound complex that is essential for mitochondrial metabolism^1, 2^. It comprises 14 catalytic core subunits that are conserved across all kingdoms of life, and up to 31 supernumerary subunits that contribute to its stability, regulation, and/or biogenesis^3–5^. As an entry point into the electron transport chain (ETC), complex I is a key contributor to oxidative phosphorylation, mitochondrial homeostasis, and redox balance. Specifically, it couples electron transfer from NADH to ubiquinone (Q) to proton translocation across the inner mitochondrial membrane, building the proton motive force (Δp) to drive ATP synthesis. Due to its central roles in metabolism, complex I is implicated in ischaemia-reperfusion (IR) injury^6^ and its dysfunctions lead to neuromuscular and metabolic diseases^7, 8^.

Two biochemically characterised, physiologically relevant states, the ‘active’ and ‘deactive’ resting states, have previously been identified by electron cryomicroscopy (cryo-EM) on mammalian complex I. They are distinguished by subtle domain movements and conformational changes at the Q-binding site and in adjacent membrane-domain subunits^5, 9–13^. In the absence of substrates and at physiological temperatures, mammalian complex I relaxes gradually from the ready-to- catalyse active resting state into the profound deactive resting state, with a partially unstructured Q-binding site^5, 9, 10, 14–17^. The deactive state forms spontaneously during ischaemia (when lack of O2 prevents ETC turnover). Then, when NADH and ubiquinone are added/replenished (for example, upon reperfusion), it slowly reactivates and returns to catalysis. During reperfusion the deactive state protects against IR injury by minimising complex I-mediated production of reactive oxygen species by reverse electron transfer^13, 18, 19^. Cryo-EM studies of mammalian complex I have also identified a third state of particles that lack key regions of density, which we suggested previously to arise from complex I in the first stages of dissociation^5, 11^.

The Q-binding site in complex I is an unusually long and heterogeneous channel. Advances in cryo-EM have allowed identification of several inhibitors^12, 20–24^, adventitious detergents^25^, and substrate analogues^21, 22, 26, 27^ bound in the site, but native Q species (Q9 and Q10, with nine and ten isoprenoid units) have so far only been observed with their Q-headgroups close to the channel entrance^28–30)^. For catalysis, the headgroup must enter much further in and approach its proposed ligating partners (His59^NDUFS2^ and Tyr108^NDUFS2^) at the tip of the channel^31, 32^ — as observed for the short-chain substrate analogues decylubiquinone (dQ)^21, 22, 27^ and piericidin A^12, 21^, and predicted by molecular dynamics simulations^33–36^. In order to position its headgroup in this reactive centre, the native substrate must span the entirety of the channel.

Here, we describe the structure of bovine complex I reconstituted into phospholipid nanodiscs with exogenous Q10. The nanodiscs provide a native membrane-like environment and eliminate potential artefacts from the detergent micelle typically used in cryo-EM analyses. Using cryo-EM, we resolve five distinct structures at global resolutions up to 2.3 Å, including, for the first time, a Q10 molecule occupying the full length of the Q-binding channel. By comparing the structures of substrate/ligand-bound and -apo (substrate/ligand-free) forms of both the active and deactive states, as well as a structure of the poorly characterised third state, we probe substrate/ligand-driven conformational changes in the Q-binding site and the physiological and catalytic relevance of each state.

## Results

### Reconstitution of bovine complex I into nanodiscs

Complex I was purified from bovine heart mitochondria in the detergent *n*-dodecyl β-D-maltoside (DDM)^10, 37^. Then, to exchange the detergent micelles for nanodiscs, it was reconstituted in a mixture of synthetic phospholipids, Q10, and the membrane scaffold protein MSP2N2^38^, using a protocol derived from that for preparing complex I proteoliposomes^23, 37, 39^. The nanodisc-bound complex I (CxI-ND) was then isolated by size-exclusion chromatography (from proteoliposomes, protein-free nanodiscs, free MSP2N2, and residual detergent) and shown to be monodisperse (Supplementary Fig. 1a). Following addition of CHAPS (3-((3-cholamidopropyl) dimethylammonio)- 1-propanesulfonate) and asolectin to dissociate the scaffold proteins and provide a larger hydrophobic phase, the sample used for cryo-EM exhibited 86.0 ± 0.1% (17.5 ± 0.4 µmol min^-1^ mg^- 1^) of the piericidin-sensitive NADH:dQ oxidoreductase activity of the DDM-solubilised enzyme before reconstitution (20.4 ± 0.6 µmol min^-1^ mg^-1^), indicating that the complex I in CxI-ND is highly catalytically competent. The intact CxI-NDs (without CHAPS/asolectin) displayed very little piericidin-sensitive NADH:dQ activity (2.01 ± 0.10 µmol min^-1^ mg^-1^) indicating that dQ does not exchange effectively in and out of the nanodisc and/or Q-binding site (Supplementary Fig. 1b).

### Resolution of three major classes of CxI-ND particles

Single-particle cryo-EM analyses on a Titan Krios microscope with a K3 detector yielded a total of 343,213 particles (Table 1), which were separated into three major classes by 3D classification in RELION^40^, following subtraction of the nanodisc density^41^ (Supplementary Fig. 2). By map-to-map comparisons with biochemically characterised mouse and bovine structures^5, 9, 10, 12, 13^ (Supplementary Table 1a), two classes were assigned to complex I in the ‘active’ resting state (61,658 particles, 2.65 Å resolution) and in the ‘deactive’ resting state (259,540 particles, 2.28 Å). The third class (22,019 particles, 3.02 Å) matched best to a state proposed earlier to correspond to particles in the first stages of dissociation; here we name it ‘state 3’ to reassess it without the bias from early suggestions based on 5.60-Å low-resolution data^5, 11^. Focussed 3D classifications subsequently resolved two sub-states for each of the active and deactive states, providing a total of five distinct maps and models (Table 1 and Supplementary Figs. 2, 3 and 4).

**Table 1.** Cryo-EM data collection, refinement, and validation statistics

|  | Data collection I |  | Data collection II |  |  |
| --- | --- | --- | --- | --- | --- |
| **Data Collection and Processing** | Grid I |  | Grid II |  |  |
| Magnification | 130,000 |  | 130,000 |  |  |
| Voltage (kV) | 300 |  | 300 |  |  |
| Electron exposure (e<sup>-</sup>/Å<sup>2</sup>) | 40.5 |  | 40.5 |  |  |
| Defocus range (μm) | -1.0 to -2.4 |  | -1.0 to -2.4 |  |  |
| Super-resolution pixel size (Å) | 0.535 |  | 0.535 |  |  |
| Final pixel size (Å) | 0.750 |  | 0.750 |  |  |
| Symmetry imposed | C1 |  | C1 |  |  |
| Initial particle images (no.) | 701,236 |  | 377,697 |  |  |
| Final particle images (no.) | 212,841 |  | 130,372 |  |  |
| Total final particle images (no.) |  |  | 343,213 |  |  |
| **Classes** | **Active** |  | **Deactive** |  | **State 3** |
|  | **Q<sub>10</sub>** | **Apo** | **Ligand<sup>1</sup>** | **Apo** | **(Slack)** |
|  | **EMD-14132** | **EMD-14133** | **EMD-14134** | **EMD-14139** | **EMD-14140** |
|  | **PDB-7QSK** | **PDB-7QSL** | **PDB-7QSM** | **PDB-7QSN** | **PDB-7QSO** |
| Final particle images (no.) | 23,449 | 38,205 | 235,957 | 23,583 | 22,019 |
| Map resolution (Å) | 2.84 | 2.76 | 2.30 | 2.81 | 3.02 |
| FSC threshold: 0.143 |  |  |  |  |  |
| Map resolution range (Å) | 2.61–7.02 | 2.49–6.70 | 2.05–3.99 | 2.48–7.16 | 2.61–6.49 |
| Map sharpening *B*-factor (Å<sup>2</sup>) | -34 | -42 | -(49–53) | -44 | -54 |
| **Model Statistics** |  |  |  |  |  |
| Initial model (PDB ID) | 7QSD | 7QSD | 7QSD | 7QSD | 7QSD |
| Model resolution (Å) | 2.85 | 2.73 | 2.23 | 2.73 | 2.91 |
| FSC threshold: 0.5 |  |  |  |  |  |
| Model composition |  |  |  |  |  |
| Non-hydrogen atoms | 69,987 | 69,743 | 71,642 | 69,322 | 66,114 |
| Protein residues | 8,287 | 8,283 | 8,259 | 8,198 | 8,015 |
| Ligands | 45 | 42 | 48 | 45 | 32 |
| Waters | 1,060 | 1,024 | 2,774 | 1,096 | 0 |
| Average *B* factors (Å<sup>2</sup>) |  |  |  |  |  |
| Protein | 49.62 | 43.88 | 40.03 | 42.27 | 51.36 |
| Ligand | 50.65 | 44.98 | 42.75 | 43.24 | 50.30 |
| Water | 44.47 | 37.52 | 37.10 | 36.74 | N/A |
| Root Mean Square deviations |  |  |  |  |  |
| Bond lengths (Å) | 0.007 | 0.009 | 0.006 | 0.006 | 0.006 |
| Bond angles (°) | 0.641 | 0.732 | 0.647 | 0.648 | 0.618 |
| MolProbity score | 1.30 | 1.36 | 1.13 | 1.37 | 1.44 |
| All-atom clash score | 3.18 | 3.76 | 2.69 | 3.68 | 4.03 |
| EMRinger score | 4.82 | 5.15 | 6.42 | 4.68 | 4.07 |
| Rotamer outliers (%) | 0.15 | 0.04 | 0.58 | 0.01 | 0.00 |
| Ramachandran plot |  |  |  |  |  |
| Favoured (%) | 96.87 | 96.83 | 97.64 | 96.69 | 96.21 |
| Allowed (%) | 3.07 | 3.13 | 2.34 | 3.28 | 3.75 |
| Outliers (%) | 0.06 | 0.04 | 0.01 | 0.02 | 0.04 |

All the established hallmarks for the active state^5, 10, 12^ are present in the active maps/models, including well-defined densities for the loops in NDUFS2, ND3 and ND1 (residues 52-62, 24-55 and 194-217, respectively) that constitute the Q-binding site, an extended NDUFA5/NDUFA10 interface, and a fully α-helical ND6-transmembrane helix (TMH) 3. [Note we use the human subunit nomenclature throughout for simplicity]. The same is true for the deactive maps/models, where the hallmarks include disordered/alternate conformations of the above loops, a restricted NDUFA5/NDUFA10 interface, and a π-bulge in ND6-TMH3^5, 9, 10, 13^. Furthermore, NDUFS7 residues 47-51 form a loop in the active state and a β-strand in the deactive state, and the adjacent loop (residues 74-83) is ‘flipped over’ between the two states, reorientating the hydroxylated conserved Arg77^NDUFS^^7^. As reported previously^5^, the state 3 map (see Supplementary Fig. 5) lacks clear density for the C-terminal half of the ND5 transverse helix and its TMH16 anchor; subunit NDUFA11 (barring a short fragment facing the intermembrane space); and the ∼40-residue N-terminus of NDUFS2. These densities remain unclear at the current (much higher) resolution, confirming these elements are disordered or in multiple unresolved conformations. Here, we focus on the occupancy of the Q-binding channel in each state and present a Supplementary Discussion to evaluate the biochemical relevance of the state 3 structure.

### Characterisation of nanodisc-bound complex I

In all three states, two belt-like densities representing two MSP2N2 monomers are visible, enveloping the membrane domain in place of the usual detergent:phospholipid micelle (Fig. 1). Two 391 residue-long polyalanine models fit into them well, overlapping as four parallel helices below NDUFA9 (Fig. 1b), a subunit just above the expected membrane surface that has been observed to bind phospholipids^10^. The overlapping contrasts with the ‘dangling’ termini observed for shorter MSPs^42^. Quantitative biochemical analyses showed the CxI-NDs contain an average of 295 phospholipids, sufficient for a layer only one or two deep around the enzyme, and 2.66 Q10. Although this equates to ∼12 mM Q10 in the phospholipid phase, substantially above the *KM* value in proteoliposomes^37^, there is no ‘bulk’ bilayer to support quinone diffusion, consistent with the low dQ/Q10 reductase activity (Supplementary Fig. 1b). Furthermore, the MSP2N2s wrap tightly around the enzyme like stretched rubber bands, creating direct enzyme-MSP2N2 contacts in some regions and phospholipid-filled crevices and cavities where there are protein voids in others (Fig. 1 and Supplementary Fig. 6a).

**Fig. 1.**
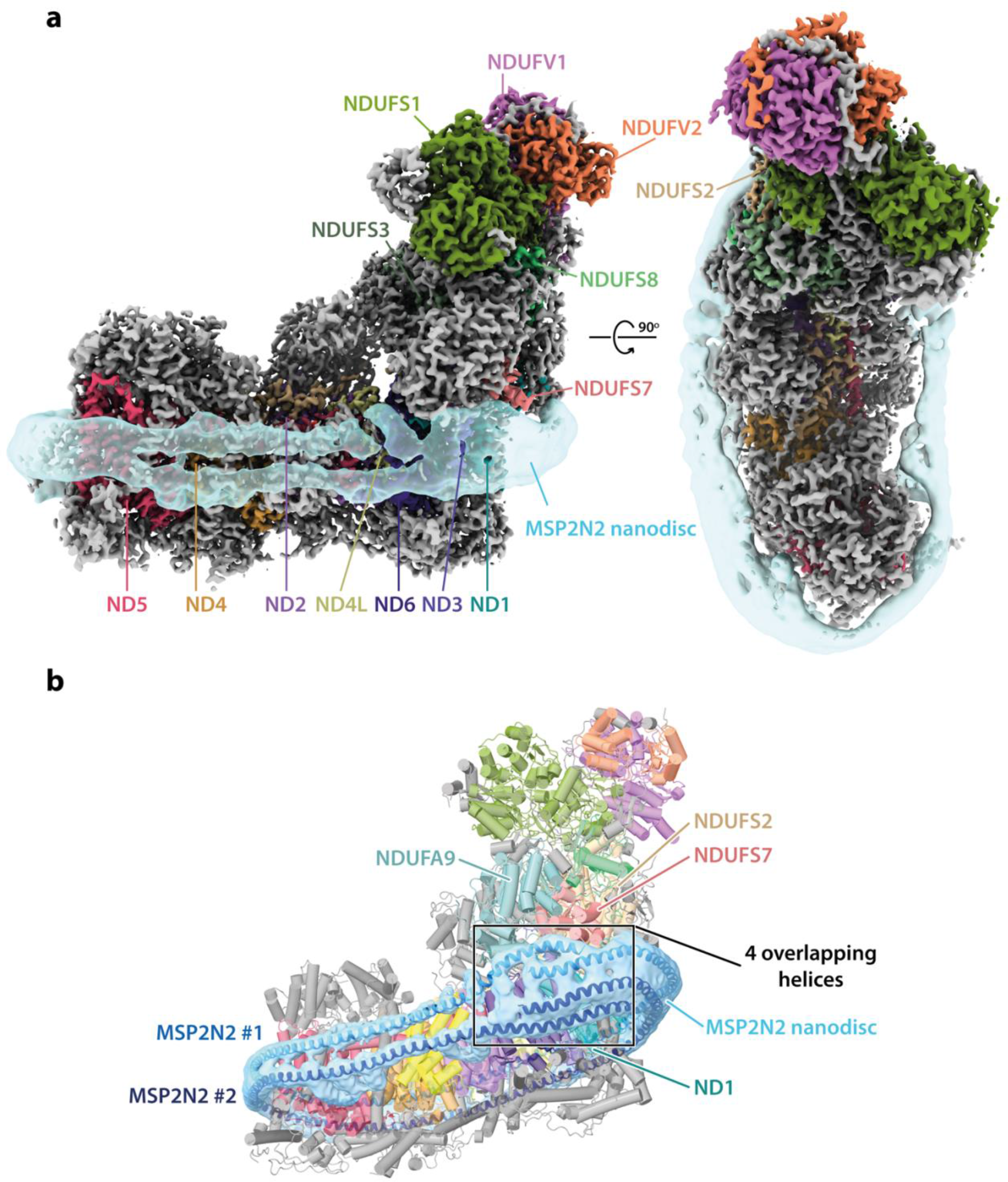
Overview of the structure of mitochondrial complex I from *Bos taurus* reconstituted into nanodiscs. **a)** Side and top views of the cryo-EM densities for the 14 core (coloured) and 31 supernumerary (grey) subunits of Q10-bound active complex I at contour level of 6.5 are shown with the subtract-refined MSP2N2 nanodisc (transparent sky blue) overlaid in UCSF ChimeraX^53^. **b)** Heel view of a representative CxI-ND model (deactive-ligand; cartoon) encapsulated in the subtract-refined MSP2N2 nanodisc map with polyalanine models for the MSP2N2 molecules shown as helices.

Comparison of the CxI-ND active-state structure with a DDM-solubilised active-state bovine structure [Protein Data Bank (PDB) ID: 7QSD; Electron Microscopy Data Bank (EMDB) ID: 14127]^43^ revealed no material differences, with convincing map-to-map correlation (0.97) and RMSD values (0.26-0.31 Å for the membrane-bound core subunits, 0.32-0.35 Å overall). As expected, none of the three DDMs observed in the reference structure^43^ were observed in CxI-NDs, while the phospholipids increased from 22 to 42 (Supplementary Fig. 6b). There is no evidence that ‘lateral pressure’ tightens the subunit interfaces in the CxI-ND membrane domain, and further comparisons with detergent-solubilised mammalian complex I structures^9, 10, 12, 13, 22, 23, 43^ revealed only three very minor structural differences (see Supplementary Discussion).

### Q10 bound in the active state

Following focussed 3D classification to resolve heterogeneous density features in the Q-binding site (see Methods and Supplementary Fig. 3), the active class was split into a ligand-bound sub- class with a continuous density matching a Q10 molecule spanning the Q-binding channel (henceforth active-Q10; 23,449 particles, 2.84 Å resolution), and a substrate-free sub-class with a presumably water-filled cavity (henceforth active-apo; 38,209 particles, 2.76 Å) (Table 1 and Supplementary Fig. 4). The Q-binding site is fully structured in both cases.

In active-Q10, the Q10 density occupies the entirety of the channel, starting between subunits NDUFS2 and NDUFS7 (with the Q-headgroup adjacent to Tyr108^NDUFS2^ and His59^NDUFS2^) then extending, as the isoprenoid chain, along the NDUFS2-NDUFS7 then ND1-NDUFS7 interfaces to exit from ND1. The His sidechain forms a hydrogen bond (H-bond) with the 3-methoxy of the Q- headgroup, rather than with either of the reactive carbonyls, which are in geometrically unfavourable positions (Fig. 2a). The Tyr sidechain is too distant (>4.3 Å) from the headgroup for a H-bond, but interacts via two mediating water molecules and a water is also H-bonded between the 4-carbonyl and Asp160^NDUFS2^. In an alternative lower-occupancy orientation of the Q10, represented by weaker density, the headgroup is flipped by 180° and the first isoprenoid is in a different position (Fig. 2a inset). While the His now interacts with the 2-methoxy the H-bonding pattern is similar. In both cases, attempts to reposition the headgroup to create reactive H-bonds to the Tyr and His without moving the isoprenoid chain out of its density were unsuccessful. A number of waters were also observed adjacent to the isoprenoid chain, stabilised by H-bonding to nearby sidechains, clustered particularly in the more charged protein section around the middle of the isoprenoid chain^37^ (Fig. 2b).

**Fig. 2.**
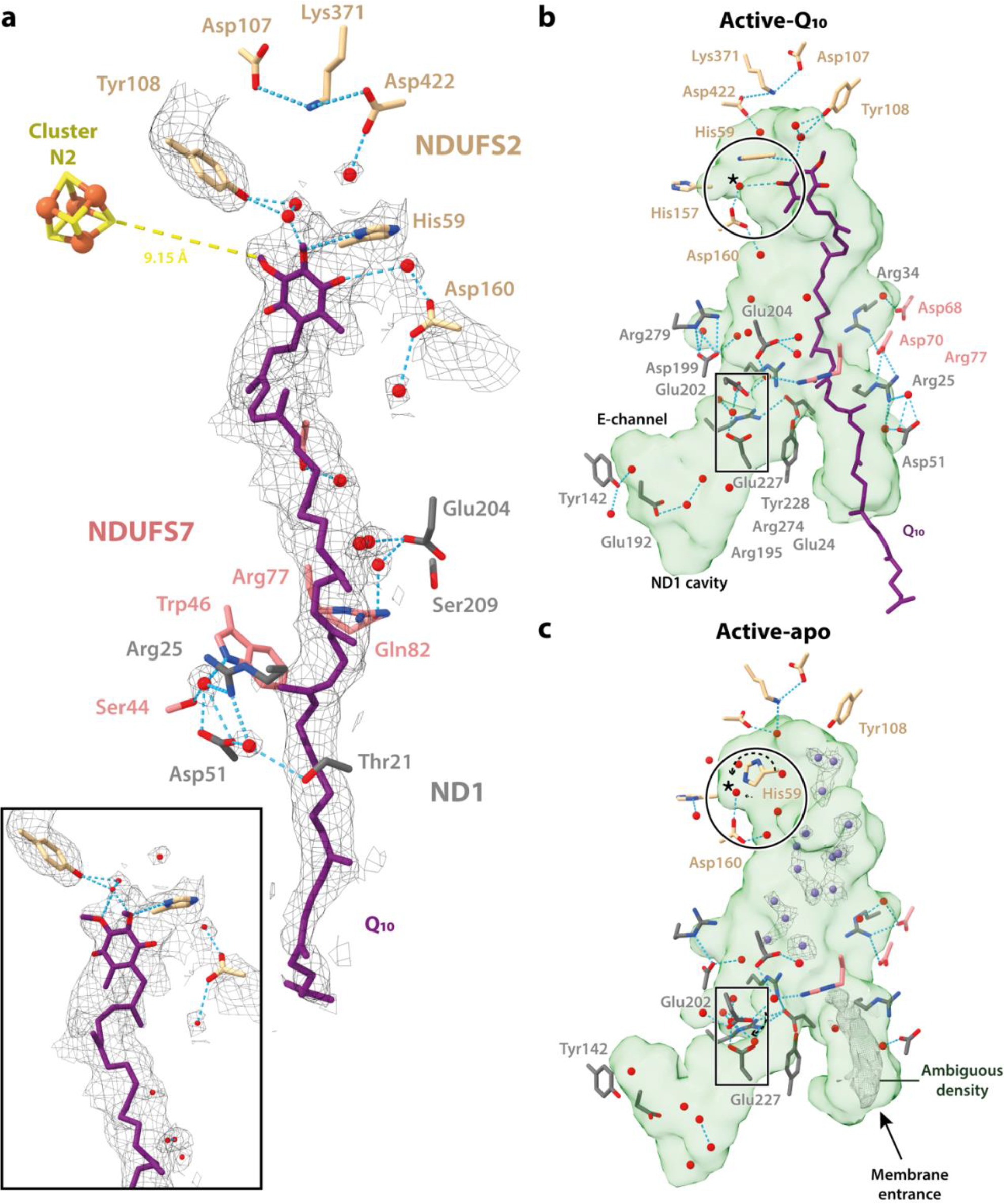
Active states of complex I with and without bound Q10. **a)** Cryo-EM densities of Q10 and neighbouring water molecules in the active-Q10 map at a contour level of 4.4 are shown together with the polar residues that make H-bonding interactions, as identified using the *hbonds* command in UCSF ChimeraX^53^. The inset shows an alternate conformation of Q10 that can be modelled into the density. The independent model-map CCmask values for the primary and secondary Q10 poses are 0.72 and 0.70, respectively. **b-c)** Protonatable residues, water molecules (red spheres) and ligands in close proximity to the Q-binding channel and ND1 cavity (as identified by CASTp^47^) are shown and labelled in the **(b)** active-Q10 and **(c)** active-apo states. Star (*) indicates a water molecule that moves as a result of the rotation of His59^NDUFS2^ (dotted arrow; circled). The unidentified density and densities for the dynamic water molecules (violet spheres) in the active-apo map are shown at a contour level of 4.0 in UCSF ChimeraX^53^.

To probe further why the Q10 appears to have ‘paused’ in this pre-reactive conformation, we applied molecular dynamics simulations with enhanced sampling to explore the charge state and flexibility of reactive groups in the Q-binding site (see Methods for details). A collective variable (CV) was used to describe the Q-headgroup position along the binding channel, with lower values denoting the headgroup closer to Tyr108^NDUFS2^ (Fig. 3a-d). Three charge-states were tested, all with cluster N2 oxidised but different protonation states for His59^NDUFS2^ and Asp160^NDUFS2^ (Figs. 2a and 3a). The free energy profile obtained for the [AspH + His] charge-state exhibits a well-defined single minimum, indicating stable binding at CV = 23.8 Å (Fig. 3a) with the Q-headgroup position matching that observed in the active-Q10 cryo-EM model at CV = 23.6 Å (Fig. 3a). The other two charge-states ([Asp^−^ + His] and [AspH + HisH^+^]) show multiple minima, with the lowest at CV ∼25 Å having the Q- headgroup distant from the reactive site. Additional structural properties (Supplementary Fig. 7) also showed significantly better agreement with the active-Q10 cryo-EM model for the [AspH + His] charge-state. The [Asp^−^ + HisH^+^] charge-state was excluded because the cryo-EM model did not suggest an ionic-pair interaction. Thus, our simulations strongly suggest the experimental structure is in the [AspH + His] charge-state and that the Q-headgroup binding pose at the top of the Q-channel is modulated by the charge-states of nearby groups.

**Figure 3:**
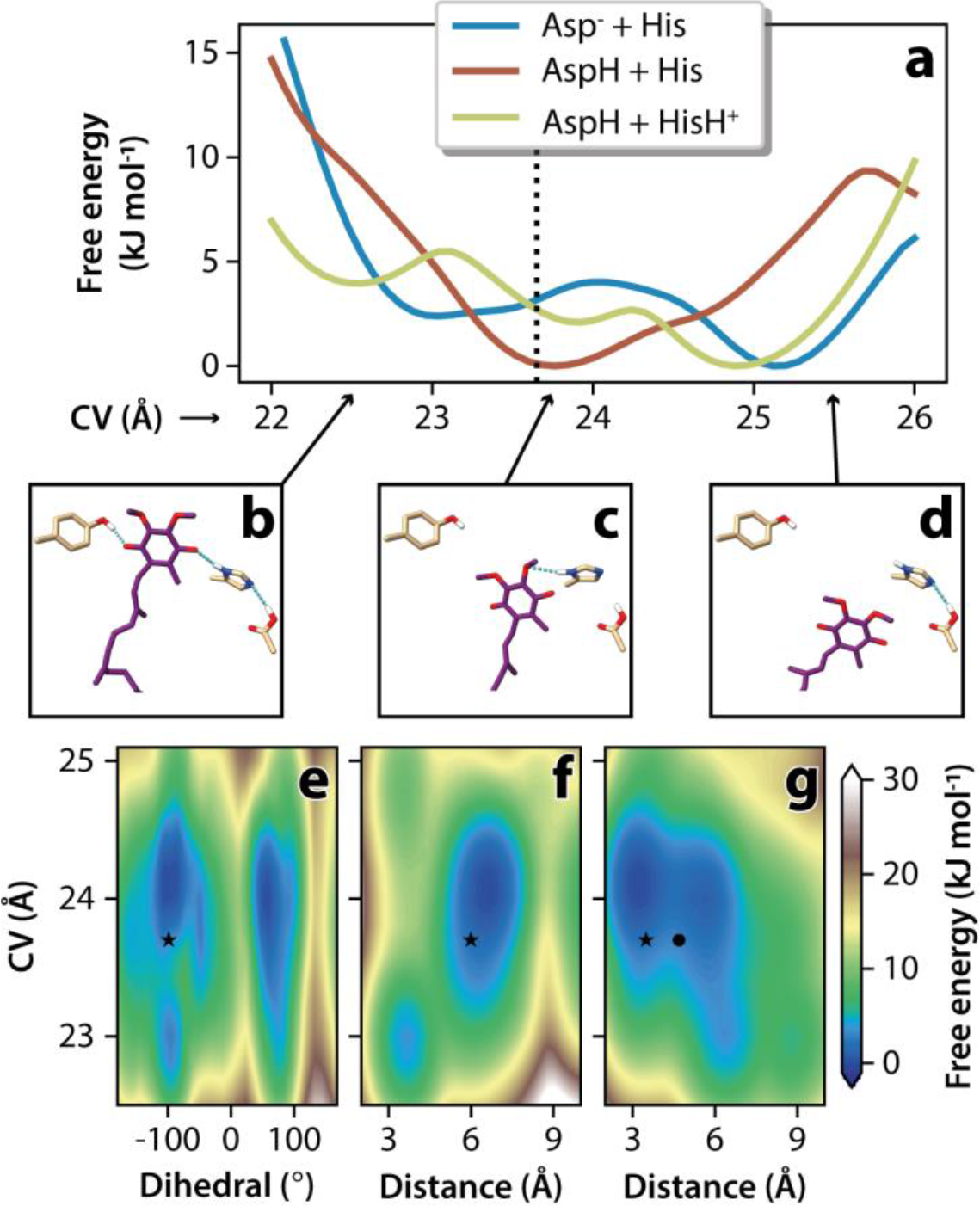
Free energy profiles from molecular dynamics simulations. **a)** Profiles were obtained for three combinations of sidechain protonation with Asp160^NDUFS2^ ionised (Asp^-^) or protonated (AspH) and His59^NDUFS2^ neutral (His, Nδ1-protonated π tautomer) or di-protonated (HisH^+^, both Nδ1- and Nε2-protonated). The collective variable (CV) describes the Q-headgroup position along the binding channel, as illustrated with His59^NDUFS2^, Tyr108^NDUFS2^ and Asp160^NDUFS2^ sidechains in **(b)** for CV = 22.5 Å, **(c)** for CV = 23.8 Å, and **(d)** for CV = 25.5 Å. The dashed line in **(a)** shows CV = 23.6 Å measured from the active-Q10 cryo-EM model. Two-dimensional profiles for the [AspH + His] charge-state along the CV coordinate and coloured by free energy are shown in **(e)** for the His59^NDUFS2^ χ2 dihedral (Cβ-Cγ bond torsion), **(f)** for the distance between the His59^NDUFS2^-Nε2 and Asp160^NDUFS2^-Cγ atoms, and **(g)** for the distance between the Q-headgroup 3-methoxy O and His59^NDUFS2^-Nδ1 atoms. Symbols correspond to the structural properties observed in cryo-EM models active-Q10 with the primary (star) and flipped (bullet) Q-headgroups.

**Fig. 4.**
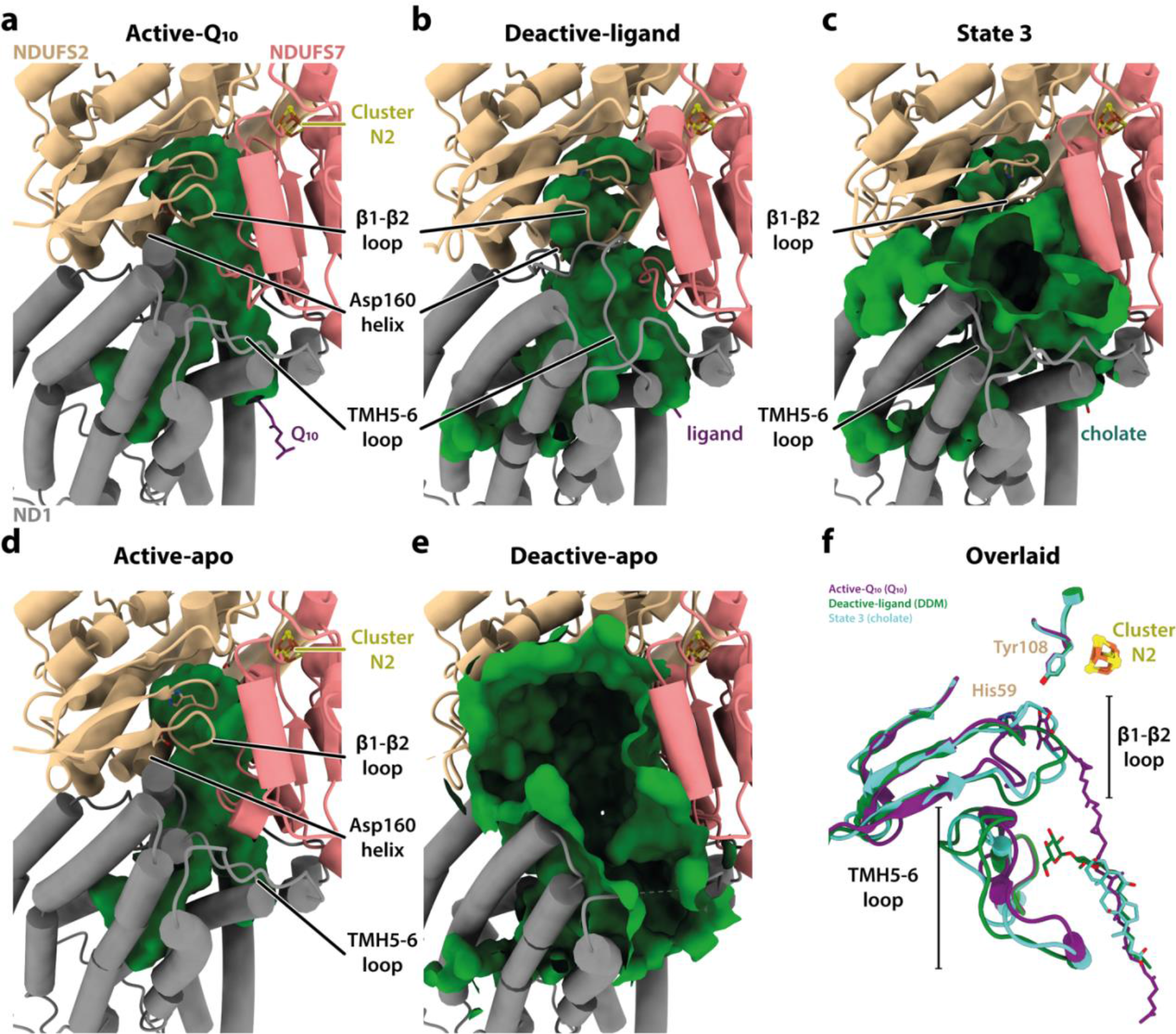
Conformations of the Q-binding site loops and the cavities they encapsulate in the five observed states of complex I. **a-e)** Cartoon representations of subunits ND1, NDUFS2, and NDUFS7 showing varying extents of disorder indicated as the (green) cavities detected by CASTp^47^. Where visible, His59^NDUFS2^, Tyr108^NDUFS2^ and Asp160^NDUFS2^ are shown as sticks. **f)** Superimposed structures of the active-Q10 (purple), deactive-ligand (green) and state 3 (turquoise) states, showing the NDUFS2-β1-β2 and ND1-TMH5-6 loops in cartoon, and His59^NDUFS2^ and Tyr108^NDUFS2^ in sticks.

Our active-Q10 model was built with the His59^NDUFS2^ dihedral angle χ2 = -95.5° (Fig. 2a), but the conformer with a ‘flipped’ sidechain also fits well. Interestingly, simulations for the [AspH + His] charge-state (Fig. 3e) revealed a flexible His59 sidechain with a barrier for the ring to flip of <15 kJ mol^-1^ at the stable binding position (CV = 23.8 Å), consistent with Cβ-Cγ bond rotamers with χ2 = - 100° or +80° being equally populated at the physiological temperature simulated. When the Q- headgroup approaches Tyr108^NDUFS2^ (CV = 22.5 Å, Fig. 3b), the flexibility of His59^NDUFS2^ χ2 is slightly decreased (Fig. 3e) and a H-bond, which is absent at the stable binding position (Fig. 3c), forms between His59^NDUFS2^-Nε2 and the protonated Asp160^NDUFS2^ sidechain (Fig. 3f). It may also re-form when the Q-headgroup dissociates (Fig. 3d), in line with previous simulations on complex I from *T. thermophilus*^34, 35^. The H-bond between His59^NDUFS2^-Nδ1 and the Q-headgroup 3-methoxy is easily broken (Fig. 3g). In fact, when the Q-headgroup occupies its stable binding position it has sufficient conformational freedom to twist and flip during the physiological-temperature simulations, also visiting configurations compatible with the flipped subpopulation observed in the cryo-EM data (Fig. 2a inset).

In the absence of contiguous density attributable to bound Q10 (active-apo structure), we suggest that the discrete densities scattered throughout the Q-binding channel, including in isoprenoid-binding regions, are dynamic water molecules in H-bonding networks (Fig. 2c). Intriguingly, there is also a clear but unidentified density observed at the channel entrance (it may be a Q10 or phospholipid inserted tail-first, but is also consistent with a molecule of MOPS buffer) that appears to separate the water-filled cavity from the hydrophobic membrane (Fig. 2c). In the reactive site, the His59^NDUFS2^ side chain is rotated by ∼90° relative to in the active-Q10 structure (Fig. 2c) and stabilised by H-bonding to the carbonyl backbone of Ile423^NDUFS2^; the water molecule between the Q-headgroup and Asp160^NDUFS2^ shifts by ∼1 Å in response (Fig. 2c). The two waters between Tyr108^NDUFS2^ and the Q-headgroup are not resolved in the active-apo sub-state, but we suggest a poorly-resolved density extending from Tyr108^NDUFS2^ represents a network of dynamic water molecules, in place of the headgroup. Apart from His59^NDUFS2^, the only difference in residue conformation between the Q-binding sites in the active-Q10 and active-apo structures is at Glu202^ND1^. Although, as may be expected for carboxylates^44^, the sidechain densities are not well-resolved, the models suggest that in active-Q10 Glu202^ND1^ forms a water-mediated H-bond to Glu227^ND1^ whereas in active-apo, Glu202^ND1^ has undergone a rotameric shift that reconfigures the H-bonding interactions between the two glutamates. These observations hint that Glu202^ND1^ responds to the channel occupancy, and may function as a ‘control point’ for proton transfer.

### Ligand binding to the deactive state

Q-binding site loops in subunits NDUFS2, ND1 and NDUFS7 are characteristically disordered in the deactive state^9, 10, 13^ and so further classification of the deactive particles focussed on the whole Q-binding region (Supplementary Fig. 3). Two sub-classes were resolved: an occupied, ligand-bound sub-class (deactive-ligand; 235,957 particles, 2.30 Å resolution) and an apparently unoccupied ‘apo’ sub-class (deactive-apo; 23,583 particles, 2.81 Å resolution) (Table 1, and Supplementary Fig. 4). In the deactive-apo structure, the Q-binding site loops in NDUFS2, ND3 and ND1 are largely disordered, whereas in the deactive-ligand structure the NDUFS2 and ND1 loops are in ordered conformations different from those in the active state (Fig. 4). The overall structure remains deactive (with a restricted NDUFA5/NDUFA10 interface, disordered ND3 loop, π-bulge in ND6- TMH3, and deactive NDUFS7 conformation). The restructured NDUFS2-β1-β2 loop (which carries His59) has moved into the space occupied by the Q-headgroup in the active-Q10 structure (Figs. 4b and f), and together with the restructured ND1-TMH5-6 loop, it constricts the Q-binding channel (Fig. 4b). In contrast, in the deactive-apo structure, the disordered loops fail to enclose the channel, which appears as a gaping crevice open to the matrix (Fig. 4e).

The ligand density observed in the deactive-ligand structure displays features consistent with both Q10 and DDM, suggesting a mixed population that could not be separated by focussed classification. The DDM used for complex I preparation may have been retained in the channel when the external DDM molecules were removed during reconstitution. The shape of the headgroup density suggests Q10 at low map thresholds but matches two six-membered maltoside rings at higher thresholds, while a long, zigzagged protrusion resembles an isoprenoid chain (Supplementary Fig. 8). Fitting a Q10 molecule into the density reveals just one polar interaction, a H-bond from Arg274^ND1^ to the Q-headgroup, while the fitted DDM molecule additionally interacts with His55^NDUFS2^ and Glu202^ND1^, and an intervening water molecule bridges it to Glu24^ND1^ (Fig. 5a). DDM may thus stabilise the deactive state, but whether it also promotes deactivation^25, 45^ remains unclear. His55^NDUFS2^ and Glu202^ND1^ are on the displaced NDUFS2-β1-β2 and ND1-TMH5-6 loops, respectively, consistent with their restructuring in the deactive-ligand structure and with continued disorder in the ND3- TMH1-2 loop, which is stabilised in the active state^25^ by interaction between His55^NDUFS2^ and Cys39^ND3^. Notably, the DDM molecule modelled here differs from the one observed in *Y. lipolytica* complex I^25^, which is slightly further into the Q-binding channel with its maltoside rings in different positions (Supplementary Fig. 9g-i). Similarly, the modelled positions of Q10 in plant complex I^29^, Q9 in *Y. lipolytica* complex I^28^, and dQ in closed and open ovine complex I^22^ (Supplementary Fig. 9m-p) overlap with the Q10/DDM modelled here, but do not align well. Clearly, the lower section of the Q-binding channel can accommodate a variety of extended hydrophobic and amphipathic molecules, including substrates, inhibitors and detergents, and their binding may modify surrounding protein structures.

**Fig. 5.**
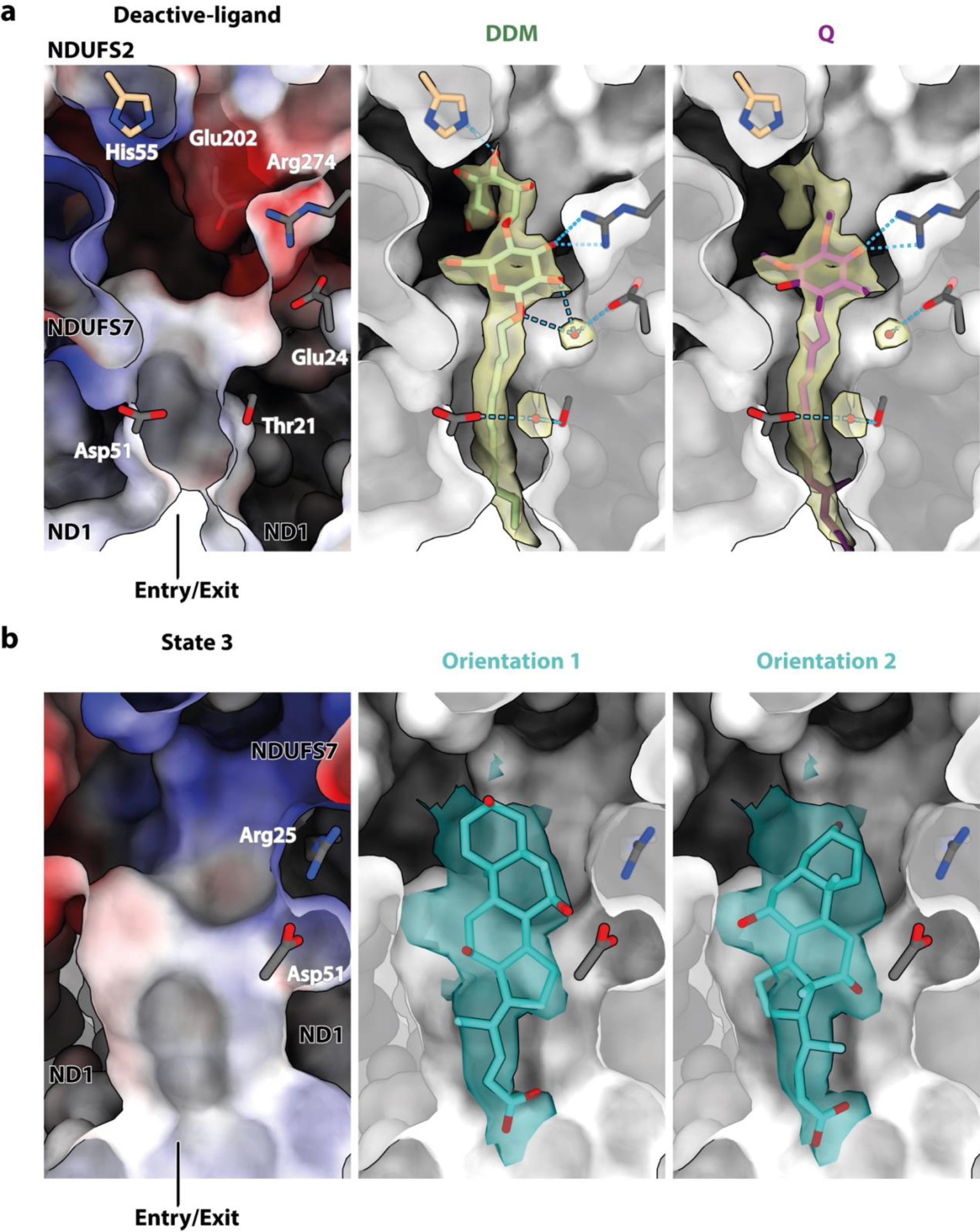
Ligands bound at the entrance of the Q-binding site of complex I. A clipped view of the surface representation of the entrance to the Q-binding channel in the **(a)** deactive-ligand and **(b)** state 3 models [coloured by Coulombic electrostatic potential (left) or by depth, in black and white (middle and right)]. **a)** Cryo-EM density of the ligand in the deactive-ligand composite map (see Methods) at a contour level of 4 (transparent yellow) in UCSF ChimeraX^53^, modelled as either a DDM (middle) or a Q10 (right). Residues H-bonded to the ligand or surrounding water molecules are shown and labelled. The model-map CCmask values for the DDM and Q10 (clipped to Q3) are 0.74 and 0.70, respectively. **b)** Cryo-EM density of the ligand in the state 3 map at a contour level of 4.5 (transparent teal), modelled as a cholate molecule in two different orientations. Residues within H-bonding distance to the ligand are shown and labelled. The model-map CCmask values for the primary and secondary cholate orientations are 0.65 and 0.67, respectively.

### Ligand binding to state 3

Following focussed classification procedures, all the state 3 particles were retained in a single homogenous class that contains densities for two bound ligands. First, a clear density at the entrance to the Q-binding channel (Fig. 4c and f) is consistent with a cholate molecule (added for the reconstitution) in two orientations (Fig. 5b), in close proximity to either Arg25^ND1^, Asp51^ND1^, and Trp46^NDUFS7^, or Arg274^ND1^, Thr21^ND1^, and Tyr228^ND1^ for polar interactions. As in the deactive- ligand structure, the NDUFS2-β1-β2 and ND1-TMH5-6 (but not ND3-TMH1-2) loops are ordered, but in different conformations to in the deactive-ligand or active states, so the shape of the Q-binding channel also differs (Fig. 4c and f). The NDUFS2-β1-β2 loop is translated up the channel relative to both its active and deactive-ligand conformations, bringing the backbone carbonyls of Ala58^NDUFS2^ and His59^NDUFS2^ to within H-bonding distance (2.6-2.8 Å) of the Tyr108^NDUFS2^ hydroxyl and restricting the cavity. The ND1-TMH5-6 loop does not run across the Q-binding channel as in the deactive-ligand state, but is retracted downwards, beyond its conformation in the active state, so the Q-binding site again appears open to the matrix. It is not possible to tell if these changes result from cholate binding or are intrinsic to state 3. Regardless of the physiological and mechanistic relevance of state 3, our structure affirms the flexible nature of the Q-binding site and its ability to accommodate ligands. Second, a clear density for an ordered Q10 molecule is observed at the extended interface between subunits ND2 and ND4 (see Supplementary Discussion and Supplementary Figs. 5a and 5b) resulting from their opposing rotation and retraction^5, 46^, and is accommodated by a π-bulge in ND4-TMH6 not present in the active or deactive states. However, the Q10 headgroup is ∼100 Å away from cluster N2 and the Q-binding channel, which is clearly too far for catalytic electron transfer.

### State-dependent structural features in the ND1 cavity and E-channel

To probe whether the different Q-site structures propagate changes to the membrane domain, we investigated the structures of the E-channel, which leads from the Q-binding site and the solvent-accessible cavity in ND1 to the first antiporter-like subunit, ND2.

The shapes of the ND1 cavities, surrounded by subunits ND1, ND3, and ND6, were visualised using the CASTp software with a 1.4 Å diameter probe^47^ (Figs. 2b-c, 4a-e and 6). In both active-Q10 and active-apo, the cavity extends past Glu227^ND1^ and Glu192^ND1^ (Fig. 2b-c) but is blocked from the next E-channel residue (Glu143^ND1^) by Tyr142^ND1^, which may thus act as a ‘proton gate’ (Fig. 6a and d). Two water molecules, on each side of Tyr142^ND1^, connect the hydrated ND1 cavity to Glu143^ND1^ and Asp66^ND3^. In both deactive-ligand and deactive-apo, the ND1 cavity extends past Glu227^ND1^, Glu192^ND1^, Glu143^ND1^, and up to Asp66^ND3^. The cavity is extended by the straightening of ND1- TMH4, which forces Tyr142^ND1^ to rotate out of the way, shifting the Glu192^ND1^ sidechain towards Glu143^ND1^ into an electrostatic interaction as described previously^22, 25, 36^, and bringing additional hydration (Fig. 6b and e). In this state, Glu143^ND1^ and Asp66^ND3^ are stabilised by two intermediary water molecules H-bonded to the backbone carbonyl of Gly62^ND6^. As in other structures in the deactive^13, 25, 27^ or open^22^ state, as well as in simulations^36^, the ND6-TMH3 π-bulge wedges Met64^ND6^ into the ∼12.5 Å-long Asp66^ND3^-to-Glu34^ND4L^ H-bond network that is continuous in the active state, disrupting it and releasing Glu34^ND4L^ to flip away towards ND2 (Fig. 6b and e). In this region, state 3 matches the deactive state (Fig. 6c). Beyond this region, we did not observe any substantial changes in the membrane domain between the active-Q10/-apo and deactive-ligand/- apo states.

**Fig. 6.**
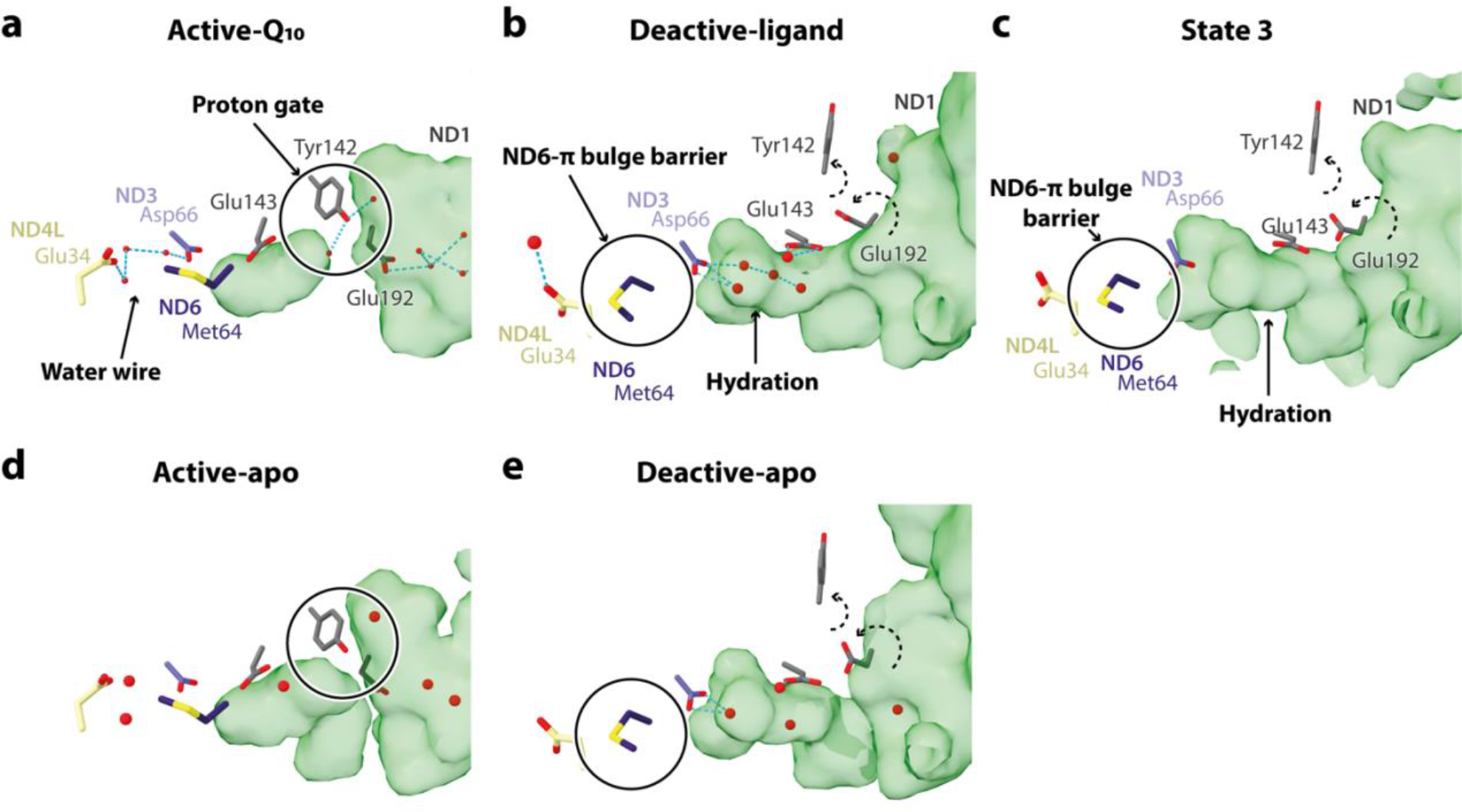
The ND1 cavity and the E-channel that connect the Q-site to the first antiporter-like subunit ND2. The proton pathways from subunits ND1 to ND4L in the **(a)** active-Q10, **(b)** deactive- ligand, **(c)** state 3, **(d)** active-apo, and **(e)** deactive-apo states. Cavities were identified by CASTp^47^ using a 1.4 Å probe. E-channel residues are shown in sticks and labelled. Dashed arrows indicate movement of residues with respect to their conformations in the active states.

## Discussion

The active-Q10 structure described here is the first structure with a native ubiquinone substrate inserted fully into the complex I Q-binding channel. However, the Q-headgroup is not bound in the expected reactive state with its 1,4-carbonyls H-bonding to Tyr108^NDUFS2^ and His59^NDUFS2^, ready to receive their protons following electron transfer from cluster N2. Simulations of the reactive state in *T. thermophilus* complex I^33, 36^ proposed that the 4-carbonyl always H-bonds to Tyr108^NDUFS2^, but the 1-carbonyl may either H-bond or π-stack with His59^NDUFS2^. Our active-Q10 structure shows none of these interactions. Furthermore, previous structures with the Q10 analogue dQ and the inhibitor piercidin A1 bound are also not modelled in the expected reactive state (Supplementary Fig. 9a-f). In the ovine ‘closed’ state supplemented with dQ and NADH^22^ the dQ-headgroup is rotated ∼35° (in-plane) relative to our primary binding pose so that His59^NDUFS2^ H-bonds to the 4-carbonyl instead of the 3-methoxy, but the 1-carbonyl is ∼4.5 Å from Tyr108^NDUFS2^. The dQ-headgroup is 5- 6 Å away from Tyr108^NDUFS2^ (Tyr144^NDUFS2^) in *Y. lipolytica* complex I also^27^. In contrast, the dQ- headgroup in a crystal structure of *T. thermophilus* complex I^21^ is flipped and rotated out-of-plane, with Tyr108^NDUFS2^ (Tyr87^Nqo4^) H-bonding with both the 4-carbonyl and 3-methoxy but His59^NDUFS2^ (His38^Nqo4^) not interacting. A similarly flipped binding pose was captured in another set of simulations on *T. thermophilus* complex I^34^, where the Q10-headgroup was stabilized by a H-bond to (protonated) His59^NDUFS2^ but did not engage Tyr108^NDUFS2^. The binding poses observed in two piericidin-bound structures^12, 21^ match the pose for dQ in the *T. thermophilus* structure^21^, although neither recapitulates the currently unique NDUFS2-β1-β2 loop conformation that increases the separation of Tyr108^NDUFS7^ and His59^NDUFS2^ in this structure.

Here, we showed excellent agreement between our experimentally determined Q10 binding pose and predictions from simulations with cluster N2 oxidised and a defined [AspH + His] charge-state for Asp160^NDUFS2^ and His59^NDUFS2^. Our results imply that the protonation and charge states of key active-site residues are intimately connected to the binding pose. Furthermore, in both our structure and simulations, cluster N2 is oxidised, so the state we observe is non-reactive for this reason also. Reduction of cluster N2 is likely a key driver for the changes required to bring the Q- headgroup into the reactive state, including ‘flipping’ the Q-headgroup to orientate the 4-carbonyl toward Tyr108^NDUFS2^, shifting local protonation equilibria to protonate His59^NDUFS2^, and changing the conformation of the NDUFS2-β1-β2 loop to bring it into a coordinating position. We suggest that our observed state is structurally equivalent to a ‘pre-reactive’ state that occurs naturally on the catalytic cycle, similar to that proposed previously by Teixeira and Arantes^35^, in which the Q10 pauses before dehydration and ligation of the 1,4-carbonyls finally brings it into the reactive conformation.

On the surface of it, our results may alternatively be taken to suggest the possibility of ubiquinone reduction in improperly ligated states, consistent with the considerable (inhibitor-sensitive) NADH:Q1/Q2 activities of the Tyr108^NDUFS2^-equivalent Y144F/W/H mutants in *Y. lipolytica*^32^. However, ubiquinone reduction potentials depend on the ligation environment and binding mode^48^, and engaging water molecules around the headgroup as non-specific proton donors appears unlikely in the context of an efficient energy-conserving mechanism. Bound Q1/Q2 have much greater conformational freedom without the long isoprenoid tail to constrain them^37, 49^, so they may exhibit alternative hydration patterns and conformational distribution^35^, and become artefactually activated for reduction.

Here, reconstituting complex I into phospholipid nanodiscs with Q10 allowed us to resolve substrate/ligand-bound and substrate/ligand-free forms of both the active and deactive states of mammalian complex I, plus a ligand-bound form of state 3. Whether deactive-like states (in which the NDUFS2-β1-β2, ND3-TMH1-2, ND1-TMH5-6 loops and a loop in subunit NDUFS7 are altered/disordered^10, 25^) are intermediates in catalysis^22^ or pronounced off-cycle resting states^14–17^ is currently a major controversy. First, we note the high similarity of our bovine/murine active and deactive states with the closed and open conformations, respectively, of ovine complex I, supported by high map-map correlations, small overall RMSD values (Supplementary Table 1b) and matching structural hallmarks^9,10, 22^. Whereas we attribute the deactive state to an off-cycle pronounced resting state, Kampjut and Sazanov^22^ proposed that ‘open’ deactive-like states must form during catalysis for Q10 to enter the Q-binding channel, which would then ‘close’ as it moves to the reactive site. Their model excludes unoccupied active-like/closed states as catalytic intermediates. However, the simplest explanation for our observation of both the active-Q10 and active-apo states, with near-identically structured Q-binding sites (as also observed in several inhibitor-bound structures^12, 22–24)^, is that Q-binding may occur within active-like/closed conformations. In this case, individual structural elements involved in the deactive transition may move during catalysis, but would not be coordinated to produce the extensive domain-level conformational changes required to generate a full deactive-like state. Simulations of Q-binding/dissociation differ in the extent to which they predict conformational changes are required^34, 35, 50–52^, and the controversy will probably only be resolved when the intermediates generated by turnover of a sample in which deactive-like states are not already present are observed. The presence of deactive-like/open states in both the pre-turnover and turnover samples characterised by Kampjut and Sazanov^22^ is consistent with both models, as they may represent off-cycle resting states that are not actively catalysing. Common to both interpretations of deactive/open states is the concept that substrate binding triggers restructuring of the Q-binding channel, either during catalysis or reactivation. Here, comparison of our deactive-ligand and deactive-apo structures shows how Q10/DDM-binding in the lower section of the channel restructures the NDUFS2-β1-β2 and ND1-TMH5-6 loops in the oxidised deactive state. These restructured loop conformations match the conformations observed in the NADH-reduced open states of ovine complex I, with ubiquinone/ubiquinol modelled in a position overlapping with that occupied here by Q10/DDM^22^. Furthermore, in the partially restructured deactive-ligand state the NDUFS2-β1-β2 loop blocks substrate access to the reactive centre, suggesting that the deactive state is resistant to reactivation under the conditions for reverse electron transport (ubiquinol and NAD^+^) because ubiquinol is unable to react with the oxidised enzyme. As a result, the deactive resting state of complex I protects against the damage caused by reverse electron transfer and the coupled generation of reactive oxygen species in ischaemia-reperfusion injury^13, 18, 19^.

Further modifications of our CxI-ND reconstitution strategy now provide new opportunities to capture and characterise hitherto-unknown catalytic intermediates that are formed as the native Q10 substrate binds and is reduced, triggering energy transfer to the proton-pumping membrane domain.

## Methods

### Transformation and recombinant expression of membrane scaffold protein, MSP2N2

Bacterial strain *Escherichia coli* NiCo21(DE3) was kindly provided by Dr Ali Ryan, Northumbria University, UK. Chemically competent *E. coli* NiCo21(DE3) cells were prepared and transformed with pMSP2N2 (Addgene) using a standard New England BioLabs (NEB) heat shock protocol. For heterologous overexpression of MSP2N2, a starter culture of *E. coli* NiCo21(DE3) containing pMSP2N2 was grown overnight at 37 °C in LB broth with 50 µg mL^-1^ kanamycin. 4 x 2 L Fernbach flasks containing 500 mL of fresh LB media supplemented with antibiotic were inoculated with 1% v/v starter culture and grown at 37 °C, 250 rpm until the culture reached an OD600 of 0.6. Protein expression was induced using 0.1 mM IPTG and the cells were cultured for a further 4 h at 37 °C with 250 rpm shaking. Cells were harvested by centrifugation at 6,000 x *g* for 20 min at 4 °C, resuspended in 40 mL lysis buffer (50 mM Tris-HCl pH 8, 500 mM NaCl, 5% glycerol, 1% v/v Triton X-100, 0.002% v/v PMSF, and 1 EDTA-free protease inhibitor tablet (Roche)) and stored at -80 °C.

### Purification of MSP2N2

The method for purification of MSP2N2 was adapted from a published protocol^38^. Cell suspensions were thawed and lysed by sonication on ice using a Q700 sonicator (Qsonica) (50% output amplitude, 30 cycles of 10 s on, 20 s off). The cell lysate was clarified by centrifugation using a SS34 rotor at 30,000 x *g* for 1 h at 4 °C. The supernatant was collected, syringe filtered through a 0.22 µm membrane (Merck Millipore Ltd.) and applied to a 1 mL Ni-NTA column (His-Trap^TM^ HP, Cytiva) equilibrated with 50 mM Tris-HCl (pH 8), 500 mM NaCl (= buffer A) + 1% v/v Triton X-100. The column was washed with 10 column volumes of buffer A + 1% v/v Triton X-100 followed by 10 column volumes of buffer A + 50 mM sodium cholate. Non-specifically bound proteins were washed with 10 column volumes of buffer A + 80 mM imidazole and finally MSP2N2 was eluted with buffer A + 400 mM imidazole. To obtain highly pure MSP2N2, the eluate from 1 mL Ni-NTA column was reapplied to a 5 mL Ni-NTA column (His-Trap^TM^ HP, Cytiva) and the same procedure was repeated. Pure MSP2N2 fractions were pooled and dialysed against 2 L of 10 mM MOPS (pH 7.5 at 4 °C), 50 mM KCl at 4 °C. Sample homogeneity was confirmed by SDS-PAGE, and the protein flash frozen and stored at -80 °C.

### Preparation of bovine mitochondria, membranes, and complex I

Bovine heart mitochondria were prepared as described previously^54^, and mitochondrial membranes prepared using a method modified from that used previously for *Mus musculus*^10^. Briefly, mitochondria were thawed and diluted to 5 mg mL^-1^ with 20 mM Tris-HCl (pH 7.55 at 20 °C), 1 mM EDTA, 10% glycerol, then ruptured by three 5 s bursts of sonication with 30 s intervals on ice using a Q700 micro-tip Sonicator (Qsonica) at 65% output amplitude setting. The membranes were pelleted at 75,000 x *g* using an MLA80 rotor (Beckman Coulter) for 1 h, then resuspended in the same buffer. Bovine complex I was prepared as described previously^37^ with a minor modification to match the mouse complex I preparation^10^ (solubilised membranes were centrifuged at 48,000 x *g* for 30 mins instead of 8,500 x *g* for 12 mins) and kept on ice until reconstitution into nanodiscs.

### Complex I reconstitution into nanodiscs

Complex I was reconstituted into nanodiscs using a protocol based on the reconstitution of complex I into proteoliposomes^23, 39^. Two batches of 0.5 mg of chloroform-dissolved synthetic lipids (1,2-dioleoyl-sn-glycero-3-phosphocholine (DOPC), 1,2-dioleoyl-*sn*-glycero-3-phosphoethanolamine (DOPE), 18:1 cardiolipin, Avanti Polar Lipids; stock: at a mass ratio of 8:1:1 (DOPC:DOPE:cardiolipin) and total concentration of 25 mg mL^-1^) were each mixed with 200 nmol of chloroform-dissolved ubiquinone-10 (Q10) in a test tube. The solvent was evaporated off under a stream of N2, and any residual chloroform removed in a desiccator under vacuum for at least 21. h. The dried lipid-Q10 mixtures were each resuspended in 457.5 µL 10 mM MOPS (pH 7.5 at 4 °C), 50 mM KCl by vigorous vortexing, and sonicated in an ultrasonic bath (Grant Instruments (Cambridge) Ltd.) for 10 min together with 42.5 µL of 20% w/v sodium cholate (i.e. final concentration of 40 mM). Each sample was transferred into a 1.5 mL Eppendorf tube, centrifuged at 7,000 x *g* for 10 min in a bench-top centrifuge, then each supernatant was transferred to a new tube and incubated on ice for 10 mins. MSP2N2 and bovine complex I (prepared as described above) were pooled gently with the lipid-Q10 mixture at a molar ratio of 400:10:1 (lipid:MSP2N2:complex I). Each sample was then diluted 2-fold with 0.5 mL 10 mM MOPS (pH 7.5 at 4 °C), 50 mM KCl to a total volume of 1 mL, and incubated on ice for 20 min. Then each sample was run over a separate PD10 desalting column (Cytiva) at 4 °C to remove the peripheral detergents. The eluates were pooled together, concentrated using a 100 kDa MWCO Amicon® Ultra concentrator (Merck Milipore Ltd.) to ∼100 µL, and filtered using a 0.22 µm Corning® Costar® Spin-X® plastic centrifuge tube filter (Merck Milipore Ltd.). The concentrated sample was applied to a Superose 6 increase 5/150 column (Cytiva) equilibrated with 10 mM MOPS (pH 7.5 at 4 °C), 50 mM KCl, and the most concentrated fractions from the monodisperse CxI-ND peak were used for grid preparation.

### Characterisation of complex I-reconstituted nanodiscs

The complex I concentration in the nanodisc preparation was quantified relative to a detergent solubilised sample of known concentration using the NADH:APAD^+^ activity assay with 500 µM APAD^+^, 1 µM piericidin and 100 µM NADH, as described previously^23, 37, 39^, except that 0.15% soy bean asolectin (Avanti Polar Lipids) and 0.15% CHAPS (Merck Chemicals Ltd.) were present. CxI-ND concentrations (i.e. combined protein concentrations of complex I and MSP2N2) were determined using the Pierce™ bicinchoninic acid (BCA) protein assay kit (Thermo Fisher Scientific). For the sample subjected to cryo-EM analyses, the complex I and CxI-ND concentrations were 3.7 and 4.8 mg mL^-1^, respectively, and the same ratio was observed consistently in several independent preparations. Phospholipid contents were determined as described previously^23, 37, 39^. The average phospholipid molecular weight was taken to be 771.6 g mol^-1^, and the volume of the phospholipid phase was estimated by assuming that 1 mg of phospholipid occupies ∼1 µL^55^. Q10 contents were quantified by HPLC, by reference to a set of standard samples, using a Nucleosil 100-5C18 column and a Dionex Ultimate 3000 RS electrochemical detector as described previously^23, 37, 39^. Q10 concentrations were defined relative to the phospholipid phase volume.

All catalytic activity assays were conducted at 32 °C in 96-well plates using a Molecular Devices Spectramax 384 plus plate reader. Catalysis was initiated by addition of 200 µM NADH and monitored at 340 and 380 nm (ε340-380 = 4.81 mM^-1^ cm^-1^). Linear rates were used for activity calculations, and inhibitor-insensitive rates (determined by the addition of 1 µM piericidin A) subtracted from each measured rate where noted. Isolated complex I used for reconstitution and CxI-ND samples used for cryo-EM analyses were diluted to 0.5 µg mL^-1^ in 20 mM Tris-HCl (pH 7.5 at 32 °C), and activity assays performed with 200 µM decylubiquinone (dQ), 0.15% asolectin/CHAPS, and/or 10 µg mL^-1^ alternative oxidase (AOX), prepared as described previously^37^.

### Cryo-EM grid preparation and image acquisition

UltrAuFoil gold grids (0.6/1, Quantifoil Micro Tools GmbH)^56^ were prepared as described previously^9^. Briefly, grids were glow discharged (20 mA, 90 s), incubated in a solution of 5 mM 11- mercaptoundecyl hexaethyleneglycol (TH 001—m11.n6-0.01, ProChimia Surfaces) in ethanol for two days in an anaerobic glovebox (Belle), then washed with ethanol and dried just before use. Using a Vitrobot Mark IV (FEI), 2.5 µL of 4.8 mg mL^-1^ CxI-ND solution (from the same preparation) were applied to the grids before blotting for 10 s at force setting -10, at 100% relative humidity and 4 °C, and then plunge-frozen into liquid ethane. Twelve grids were screened for particle number and distribution and two grids were selected for two 2-day data collections. They were imaged using a Gatan K3 detector and a post-column imaging energy filter (Gatan BioContinuum) operating in zero-energy mode with a slit width of 20 eV mounted on an FEI 300 keV Titan Krios microscope (Thermo Fisher Scientific) with a 100 µm and 70 µm objective and C2 apertures, respectively, and EPU v. 2.7.0.5806REL at the Department of Biochemistry, University of Cambridge. Data were collected in super-resolution electron counting mode at a pixel size of 0.535 Å pixel^-1^ (81,000x nominal magnification) with a defocus range -1.0 to -2.4 in 0.2 µm intervals, and the autofocus routine run every 10 µm. Aberration-free image shift (AFIS) was used for data acquisition on day 1 of the 2-day data collection for the first grid but was abandoned due to frequent occurrences of erratic image beam shifts observed in collected movies. The dose rates for the two datasets were 16.9 electrons Å^-2^ s^-1^, with 2.4 sec exposures captured in 40 frames. The total dose was thus 40.5 electrons Å^-2^ in both cases. Data were retrieved as non-gain-corrected LZW-compressed tiff movie stacks.

### Cryo-EM data processing

The two datasets were processed separately until stated otherwise, using RELION 3.1.0^40^ (Supplementary Figs. 2 and 3). The micrographs were motion-corrected using RELION’s implementation of MotionCorr with 5 x 5 patches and amplitude contrast of 0.1, and contrast transfer function (CTF) estimated using CTFFIND-4.1^57^ with ResMax set to 5 Å. Micrographs were filtered to remove those with a negative rlnCtfFigureOfMerit value, an rlnMaxResolution value worse than 6 Å, or an rlnCtfAstigmatism value lower than 20 or greater than 1000. Ice-contaminated micrographs were further removed manually to give 2,639 and 1,797 micrographs for datasets 1 and 2, respectively, from which 804,367 and 382,037 particles were selected using RELION’s AutoPicking tool with a 3D map input^43^. Particles were extracted with an initial 4.5x downscaling to 2.4075 Å pixel^-1^, and filtered to select those with an rlnAutopickFigureOfMerit value between 0 and 4. Following one round of 2D (with alignment) and 3D (without alignment) classification to remove junk particles, the remaining particles were re-extracted at the nominal pixel size (1.07 Å pixel^-1^; 2x bin) for another round of 3D classification with local angular search to remove aberrant classes of particles. 251,045 (dataset 1) and 140,731 (dataset 2) particles were brought forward for iterative rounds of CTF refinement^40, 58^, to estimate anisotropic magnification, beam tilt, trefoil, 4^th^ order aberration, and per-particle defocus, astigmatism and B-factor parameters. Particles with an rlnNrOfSignificantSamples value greater than 2,999 were removed, and the two datasets combined to give 367,615 particles. These particles were then re-extracted with re-centring, subjected to Bayesian polishing, CTF refined and 3D classified (local angular search) to remove any remaining junk. At this early stage, 358,326 particles were 3D refined with solvent flattening to give a global resolution of 2.28 Å (according to a gold-standard Fourier shell correlation (FSC) of 0.143). Signal subtraction was performed to remove most of the non-complex I contribution: a tight complex I mask was first generated from a working model using RELION MaskCreate, and subtracted from the consensus map to obtain densities for the nanodisc; leftover complex I densities from subtraction were removed using a volume eraser in UCSF Chimera^59^, and the complex I-subtracted map then used to make a mask for the nanodisc in RELION. 3D classification (number of classes, *K* = 6, local angular search to 0.2° sampling) was then performed, which separated the particles into active, deactive and state 3 complex I classes. Two classes arising from atypically shaped nanodisc complexes (C1) and junk (C2) were excluded from subsequent rounds of processing. Active (C4), deactive (C5 and C6) and state 3 (C3) classes retained 61,654, 259,547, and 22,019 particles, respectively, which were then signal reverted to include the nanodisc densities, and repolished at 0.7523 Å pixel^-1^. The three classes refined to 2.65, 2.28, and 3.02 Å resolution, respectively, at the calibrated pixel size of 0.7496 Å pixel^-1^ (Supplementary Fig. 1. 2) determined by comparison with existing mammalian complex I structures^12^. Focussed 3D classification without alignment (regularisation parameter, *T* = 100) was performed on the active, deactive, and state 3 classes (Table 1 and Supplementary Fig. 3). All individual classes were first subject to signal subtraction to retain roughly only the peripheral arm of complex I, and then focussed classified using a mask generated from a tentatively modelled Q10 (active), a mask generated from a provisional partial protein model (ND1, NDUFS2, and NDUFS7) encapsulating the Q-binding site (deactive), or a mask generated from a tentatively modelled DDM (state 3) (Supplementary Fig. 3); 7 junk particles were discarded in this step for the deactive class. Masks for focussed classification were generated using RELION MaskCreate with up to 10 pixels of extensions and soft cosine edges. The identified sub-states outlined in the main text were then signal reverted to give the global map, and the global resolution estimated from the FSC between two independent, unfiltered half-maps (FSC = 0.143) (Table 1 and Supplementary Fig. 3 and 4). As no obvious differences were identified between the state 3 sub-states, which were evenly balanced, they were kept as a single class. The model-generated mask used for 3D refinement with solvent flattening and resolution estimation was generated in UCSF Chimera^59^ using the ‘molmap’ function, before being low-pass filtered to 15 Å and having a 6-pixel soft cosine edge added using RELION MaskCreate. Mollweide projections were plotted using Python and Matplotlib.

All consensus maps were locally sharpened from the unsharpened, unfiltered half-maps generated from RELION post-process (user-provided B-factor and ad-hoc low-pass filter set to 0 and Nyquist, respectively; the output pixel size was altered to match the calibrated pixel size) using phenix.autosharpen in Phenix 1.18.2-3874^60^, setting the resolution limit to the highest local resolution determined from RELION LocalRes (Supplementary Fig. 4), and with a local sharpening box size of 15^3^ pixels and a targeted overlap of 5 pixels. The deactive-ligand map was split into three sections (distal and proximal membrane domains, and peripheral domain) for manual multibody refinement (i.e. signal subtraction, followed by focussed refinements) following nanodisc subtraction (Supplementary Fig. 3). The focus-refined maps were then globally sharpened in RELION post-process, and using the globally sharpened consensus deactive-ligand map as a reference, combined to make a composite map using phenix.combine_focused_maps in Phenix 1.19-4092^60^ (Supplementary Fig. 3). The composite map was carefully compared to the consensus map to ensure that there were no map distortions or anomalies. The locally sharpened global maps (active-Q10, active-apo, deactive-apo, and state 3) and globally sharpened composite map (deactive-ligand) were used for model building and refinement (Table 1). Nanodisc maps were generated by complex I subtraction and focussed refinement (i.e. subtract-refinement) using a nanodisc mask, either with alignment (all particles combined) or without alignment (individual sub- states).

### Model building, refinement and validation

Working models derived from a model for bovine complex I in the active state (PDB ID: 7QSD)^43^ were rigid-body fitted into maps using the *Fit in Map*tool in UCSF Chimera^59^, rigid-body real space refinement in Phenix 1.18.2-3874^60^, and Curlew all-atom-refined using Coot 0.9.4-pre^61^. The models were checked, and new resolvable regions built manually in Coot 0.9.4.2-pre. Local bovine populations are known to have a polymorphism at residue position 255 of subunit NDUFA10 – cDNA sequencing has shown evidence for both asparagine^62^ and lysine^3^, while electrospray ionisation mass spectrometry supports the latter^63^. On the basis of these reports and the Coulomb potential densities in the CxI-ND maps, we modelled it as Lys255^NDUFA^^10^. Similarly, residue position 129 (glutamine) of subunit NDUFS2 was modelled as Arg129^NDUFS2^. Densities for existing and additional phospholipid molecules were identified with the *Unmodelled blobs* tool in Coot. All DDM molecules in the starting model were removed or replaced with phospholipid molecules where the CxI-ND map features indicated, and lipid tails were clipped where necessary using the delete tools in Coot and PyMOL. The manually inspected models were then real-space refined against the respective locally sharpened consensus (active-Q10, active-apo, deactive-apo, and state 3) or composite (deactive-ligand) maps in Phenix 1.18.2-3874^60^ with Ramachandran restraints set to Oldfield (favoured) and Emsley8k (allowed and outlier) to remove genuine forced twists. This real-space refinement step was performed iteratively with manual adjustments in Coot. Water molecules were placed into distinct density peaks as identified with the *Find Waters* function in Coot, with the minimum and maximum distance to protein atoms set to 2.4 and 3.4 Å, respectively. The identified waters were manually edited to remove falsely placed waters (based on H-bonding geometries, strength and shape of densities, and steric clashes) and bulk solvent waters, and to add waters missed due to uncertain positions of surrounding side chains or waters. Atom resolvabilities (Q-scores) in the respective cryo-EM maps were calculated using MapQ^64^, and any outliers identified and corrected. The models were then real-space refined in Phenix as outlined above. The model statistics for the five sub-states in active, deactive and state 3 classes (Table 1) were produced by Phenix, MolProbity, and EMRinger. Model-to-map FSC curves were generated using phenix.validation_cryoem. Model-map CCmask values for various substrate/ligand poses were calculated by first extracting the ligand coordinates from the protein models and then running phenix.validation_cryoem against their respective final maps.

A provisional poly-alanine model for the two MSP2N2 nanodisc belts was built and refined into a 7.4 Å subtract-refined nanodisc map made from all three major species of complex I-reconstituted nanodiscs using interactive molecular dynamic simulations in ISOLDE 1.2.2^65^ with α-helical secondary structure restraints.

### Comparisons of cryo-EM maps and models

Map-to-map real-space correlations were performed using the *Fit in Map* function in UCSF ChimeraX^53^ (Supplementary Table 1) following low-pass filtering of the relevant maps to the resolution of the lowest resolution map in the set in RELION^66^. RMSD calculations were performed using the *Align* command in PyMOL^67^.

### Identification of hydrogen bonds

H-bonding contacts within individual CxI-ND models (with hydrogens added using *phenix.ready_set* and/or *phenix.reduce*^60^) were identified using the *hbonds* command in UCSF ChimeraX^53^, for which the geometric criteria are based on a survey of small-molecule crystal structures^68^, and atom types adapted and extended from the program IDATM^69^.

### Quinone cavity determination

The interior surface of the Q-binding channel was predicted using CASTp^47^, which computes a protein surface topology from a PDB model. The default 1.4 Å radius probe was used and the results were visualised in PyMOL^67^ using the CASTpyMOL 3.1 plugin and by UCSF ChimeraX^53^.

### Molecular dynamics simulations

The cryo-EM structure of active-state complex I from *Mus musculus* at 3.1 Å resolution (PDB ID: 6ZR2)^12^ was used to build the initial simulation model. Protonation states of sidechains were adjusted to neutral pH, except that His59^NDUFS2^ and Asp160^NDUFS2^ were modelled initially as di-protonated (HisH^+^) and neutral (AspH), Glu68^ND3^, Glu36^NDUFS5^, Glu262^ND1^ and Glu114^ND4^ as neutral (GluH), and His549^NDUFS1^ and His42^NDUFB2^ as di-protonated. The N-termini of NDUFS7 (Ser34) and the truncated NDUFB6 (Ser66) were modelled as neutral. These alternative protonation states were suggested by the chemical environment of the group and PropKa calculations^70^ (with nearby FeS cluster N2 in the oxidised state). All cofactors and post-translational modifications present in PDB 6ZR2 were included. High confidence phospholipids were retained and built with linoleoyl (18:2) acyl chains. The polar headgroups were preserved, thereby they were modelled as 1,2- dilinoleoyl-sn-glycero-3-phosphatidylcholine (DLPC), 1,2-dilinoleoyl-sn-glycero-3- phosphatidylethanolamine (DLPE) and 1’-3’-bis[1,2-dilinoleoyl-sn-glycero-3-phospho]-sn-glycerol, as the cardiolipin dianion (CDL). Missing hydrogen atoms were built with the GROMACS suite^71^. Neutral His tautomers were chosen based on optimal H-bonding.

The modelled protein complex was embedded^72^ in a solvated bilayer with a composition mimicking that of the inner mitochondrial membrane^73^. The final solvated and inserted system contains 368 DNPC (179 in the matrix leaflet), 294 DNPE (143 in the matrix leaflet), 96 CDL (half in each leaflet) and 22 oxidised Q10 molecules, all initially in the membrane phase. The asymmetry in DLPC and DLPE composition between the two leaflets is due to chain NDUFA9 occupying an area of the matrix leaflet only. The water phase contains 205,387 molecules plus 553 Na^+^ and 348 Cl^−^ ions to neutralize the total system charge and maintain ∼0.1 M salt concentration. The resulting simulation model contains a total of 861,975 atoms. Water solvation of the apo Q-binding channel and regions near the oxidised N2 FeS centre was compared to the active-Q10 and previous cryo-EM models (PDB ID: 6YJ4)^25^ and adjusted accordingly.

The simulation model was relaxed and equilibrated during molecular dynamics (MD) simulations of 740 ns in total. Initially all protein heavy atoms were tethered to their initial position by harmonic restraints, then the force constants were decreased progressively from 1,000 to 10 kJ mol^-1^ nm^-1^, and all atoms were free to move in the last 210 ns. Membrane packing (area per lipid) and hydration of several groups in the Q-binding channel were monitored and checked for stability after 400 ns. Then, the final configuration of this simulation had one Q10 molecule inserted into the binding channel. The Q10 coordinates, as well as those of the sidechains of His59^NDUFS2^, Tyr108^NDUFS2^,

Thr156^NDUFS2^, Met70^NDUFS7^ and Ser205^ND1^, were adjusted to the superimposed active-Q10 cryo-EM model (Fig. 2a). Clashing water molecules in the channel were removed and the protonation states of His59^NDUFS2^ and Asp160^NDUFS2^ were changed to the three charge-states studied ([Asp^-^ + His]; [AspH + His]; [AspH + HisH^+^]). If necessary, charge neutrality was maintained by removing a counter-ion. Each charge-state was further relaxed and equilibrated during 235 ns of MD simulation, with initial harmonic restraints in heavy protein atoms, the distances between atoms His59^NDUFS2^-Nδ1–Q10O3 and His59^NDUFS2^-Nε2–Asp160^NDUFS2^-Cγ, and the collective variable (CV) described below. Restraint forces were progressively reduced to zero and removed fully in the last 50 ns. Hydration of the Q-binding channel was again monitored and checked for stability. This procedure was designed and implemented to generate equilibrated and unbiased simulation models in the three charge-states studied. A canonical MD trajectory of 300 ns without any restraints was obtained for each charge-state. The root-mean squared deviation (RMSD) for the positions of the Cα atoms of chains NDUFS7, NDUFS2 and ND1 remained stable during these trajectories at ∼1.4 Å in relation to both the initial model (PDB 6ZR2) and the current active-Q10 cryo-EM model.

A pathway collective variable (CV)^74^, as used previously for Q-binding simulations^35^, was applied to describe the position of the Q-headgroup along the channel. This CV (see Fig. 3b-d) is a combination of distances between the heavy atoms in the Q-headgroup and the Cα atoms of residues in subunits NDUFS7, NDUFS2 and ND1 exposed to the Q-binding channel. Distances are evaluated with respect to four milestone configurations that represent progressive binding of Q10.

Finally, well-tempered metadynamics^75^ simulations were performed for each charge-state, starting from a configuration taken at 75 ns of each canonical MD trajectory described above. Metadynamics were activated in the CV coordinate (position on the path, p1.sss, and distance from the path, p1.zzz) and in the His59^NDUFS2^ χ2 dihedral (Cβ-Cγ bond torsion) with Gaussians deposited every 500 time steps (1 ps), at initial height of 0.6 kJ mol^-1^, widths of 0.4 and 0.02 units for the CV and dihedral, respectively, and a bias factor of 15.0. Walls were included to restrict sampling for the CV at 21 < p1.sss < 26 Å and -2.45 < p1.zzz < -2.20 Å, with force constant of 1000 kJ mol^-1^ nm^-1^. Productive metadynamics simulations lasted 140 ns for each charge-state. Due to the enhanced sampling nature of this method, simulation times significantly shorter than for canonical MD are sufficient for an appropriate conformational sampling of restricted regions, such as the reactive position of the complex I Q-binding site. Convergence within ±1 kJ mol^-1^ of free energy differences in the CV profile (Fig. 3a) was reached after 70 ns. The effects of metadynamics and of restraints were removed by re-weighting the distribution of structural properties (distances, dihedral, CV) and the resulting free energies are shown in Fig. 3a, e-g and Supplementary Fig. 7.

In all MD simulations the interactions of protein, lipids and ions were described with the all-atom CHARMM36m force-field^76–78^. Water was represented by the standard TIP3P model^79^. FeS centres were described using the Chang & Kim^80^ parameters with corrections by McCullagh & Voth^81^. Q10 interactions were represented by our calibrated force-field^73, 82^. The remaining cofactors were described by available CHARMM and CGenFF parameters (charmm36-mar2019.ff)^78^. All simulations were conducted with GROMACS (version 2020.3)^71^ at constant temperature of 310 K, pressure of 1 atm and a time step of 2 fs. Long-range electrostatics were treated with the Particle Mesh Ewald method^83^. Metadynamics simulations were performed with the PLUMED plugin (version 2.6.1)^84^.

### Data availability statement

Data supporting the findings of this manuscript, and simulation workflow scripts, are available upon reasonable request. Data accession codes: EMD-14132 and PDB ID: 7QSK (active-Q10), EMD- 14133 and PDB ID: 7QSL (active-apo), EMD-14134 and PDB ID: 7QSM (deactive-ligand; composite), EMD-14135, EMD-14136, EMD-14137, and EMD-14138 (deactive-ligand; consensus, hydrophilic domain, proximal and distal membrane domains, respectively), EMD-14139 and PDB ID: 7QSN (deactive-apo), and EMD-14140 and PDB ID: 7QSO (state 3).

## Acknowledgements

We thank D. Chirgadze, S. W. Hardwick and L. Cooper (University of Cambridge Cryo-EM facility) for assistance with grid screening and cryo-EM data collection during the COVID-19 pandemic with restricted access; A. Ryan (Northumbria University, UK) for the *E. coli* NiCo21 (DE3) expression strain for MSP2N2; the SDumont cluster in the National Laboratory for Scientific Computing (LNCC/MCTI, Brazil) for computational resources; and D. N. Grba (MRC MBU) for critical feedback. This work was supported by the Medical Research Council (MC_UU_00015/2 to J.H.), the Swiss National Science Foundation (P400PB_191096 to O.B.) and Fundação de Amparo à Pesquisa do Estado de São Paulo (FAPESP, grant 2019/21856-7 to G.M.A. and fellowship 2020/14542-3 to C.S.P.).

## Author contributions

I.C. performed CxI-ND reconstitution, cryo-EM grid preparation, collected and processed cryo-EM data, carried out structure model building, analyses, and interpretations, and prepared figures. I.C. and J.J.W. carried out biochemical and kinetic characterisations. J.J.W. prepared complex I for reconstitution, assisted by I.C. H.R.B. advised on cryo-EM data collection and processing and built initial models. J.J.W. and H.R.B. contributed to data interpretation. B.S.I. prepared MSP2N2 for reconstitution, with contributions from J.J.W. O.B. established and optimised the initial reconstitution protocol. C.S.P. and G.M.A. performed and analysed molecular simulations. J.H. initiated and supervised the project and contributed to interpretation of the data. I.C. and J.H. wrote the paper with input from all authors.

## SUPPLEMENTARY FIGURES & TABLES

**Supplementary Table 1:**
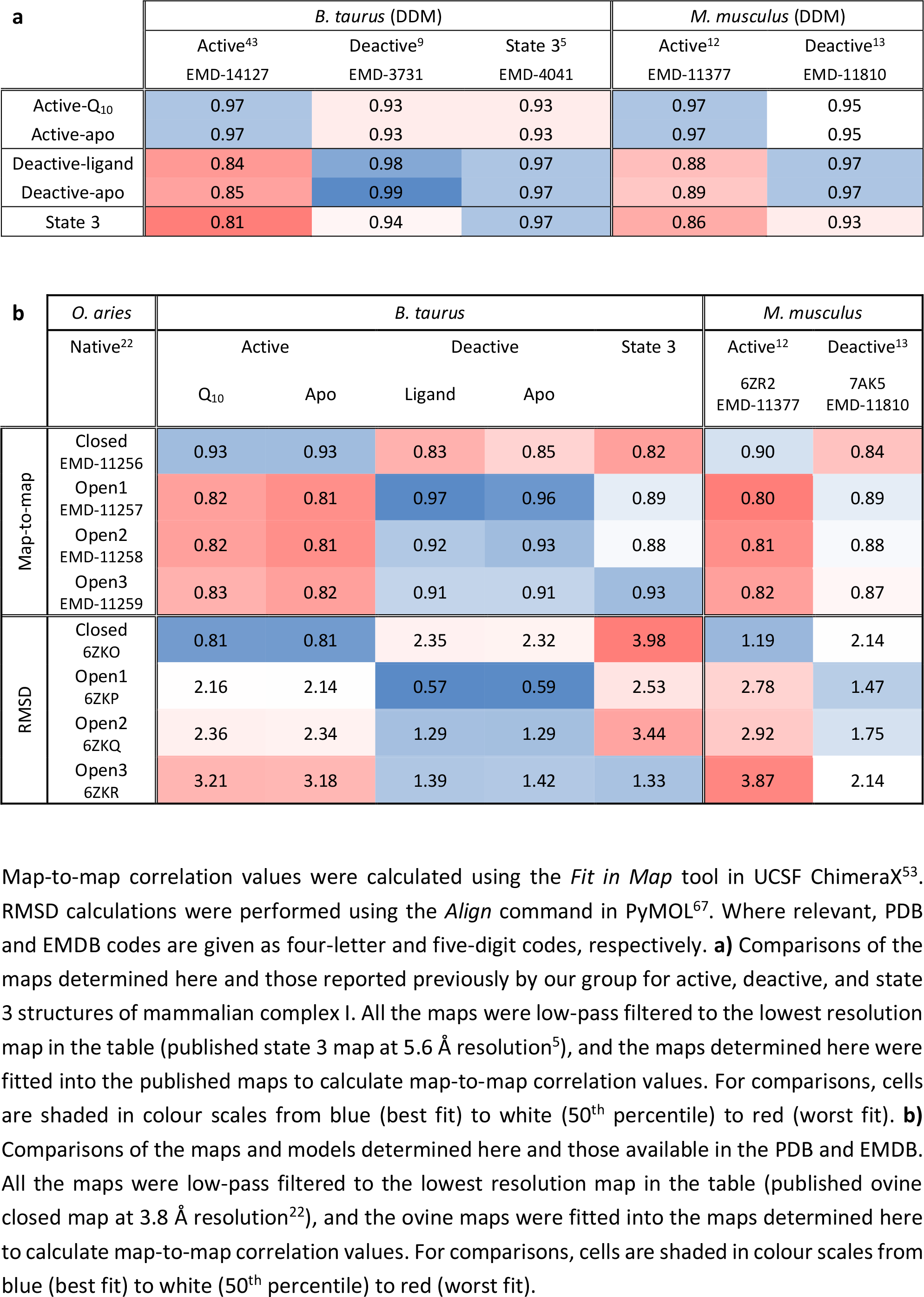
Map-to-map correlations and RMSD values between maps and models determined here and available in the protein and electron microscopy data banks.

| **a** | *B. taurus* (DDM) |  |  | *M. musculus* (DDM) |  |
| --- | --- | --- | --- | --- | --- |
|  | Active<sup>43</sup> | Deactive<sup>9</sup> | State 3<sup>5</sup> | Active<sup>12</sup> | Deactive<sup>13</sup> |
|  | EMD-14127 | EMD-3731 | EMD-4041 | EMD-11377 | EMD-11810 |
| Active-Q<sub>10</sub> | 0.97 | 0.93 | 0.93 | 0.97 | 0.95 |
| Active-apo | 0.97 | 0.93 | 0.93 | 0.97 | 0.95 |
| Deactive-ligand | 0.84 | 0.98 | 0.97 | 0.88 | 0.97 |
| Deactive-apo | 0.85 | 0.99 | 0.97 | 0.89 | 0.97 |
| State 3 | 0.81 | 0.94 | 0.97 | 0.86 | 0.93 |

| b | *O. aries* | *B. taurus* |  |  |  |  | *M. musculus* |  |
| --- | --- | --- | --- | --- | --- | --- | --- | --- |
|  | Native<sup>22</sup> | Active |  | Deactive |  | State 3 | Active<sup>12</sup> | Deactive<sup>13</sup> |
|  |  | Q<sub>10</sub> | Apo | Ligand | Apo |  | 6ZR2<br>EMD-11377 | 7AK5<br>EMD-11810 |
| Map-to-map | Closed<br>EMD-11256 | 0.93 | 0.93 | 0.83 | 0.85 | 0.82 | 0.90 | 0.84 |
|  | Open1<br>EMD-11257 | 0.82 | 0.81 | 0.97 | 0.96 | 0.89 | 0.80 | 0.89 |
|  | Open2<br>EMD-11258 | 0.82 | 0.81 | 0.92 | 0.93 | 0.88 | 0.81 | 0.88 |
|  | Open3<br>EMD-11259 | 0.83 | 0.82 | 0.91 | 0.91 | 0.93 | 0.82 | 0.87 |
| RMSD | Closed<br>6ZKO | 0.81 | 0.81 | 2.35 | 2.32 | 3.98 | 1.19 | 2.14 |
|  | Open1<br>6ZKP | 2.16 | 2.14 | 0.57 | 0.59 | 2.53 | 2.78 | 1.47 |
|  | Open2<br>6ZKQ | 2.36 | 2.34 | 1.29 | 1.29 | 3.44 | 2.92 | 1.75 |
|  | Open3<br>6ZKR | 3.21 | 3.18 | 1.39 | 1.42 | 1.33 | 3.87 | 2.14 |

Map-to-map correlation values were calculated using the *Fit in Map* tool in UCSF ChimeraX<sup>53</sup>. RMSD calculations were performed using the *Align* command in PyMOL<sup>67</sup>. Where relevant, PDB and EMDB codes are given as four-letter and five-digit codes, respectively. **a)** Comparisons of the maps determined here and those reported previously by our group for active, deactive, and state 3 structures of mammalian complex I. All the maps were low-pass filtered to the lowest resolution map in the table (published state 3 map at 5.6 Å resolution<sup>5</sup>), and the maps determined here were fitted into the published maps to calculate map-to-map correlation values. For comparisons, cells are shaded in colour scales from blue (best fit) to white (50<sup>th</sup> percentile) to red (worst fit). **b)** Comparisons of the maps and models determined here and those available in the PDB and EMDB. All the maps were low-pass filtered to the lowest resolution map in the table (published ovine closed map at 3.8 Å resolution<sup>22</sup>), and the ovine maps were fitted into the maps determined here to calculate map-to-map correlation values. For comparisons, cells are shaded in colour scales from blue (best fit) to white (50<sup>th</sup> percentile) to red (worst fit).

**Supplementary Fig. 1:**
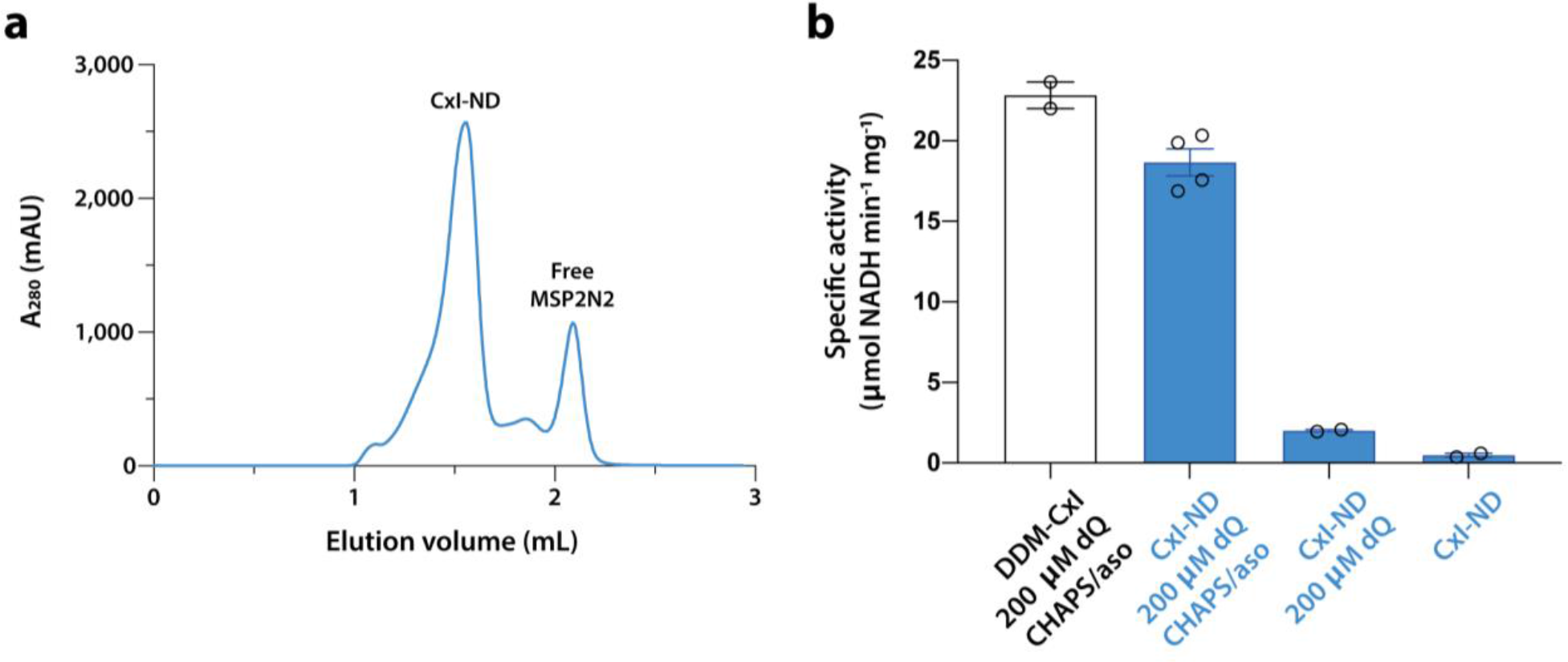
Purification and biochemical characterisation of *Bos taurus* complex I reconstituted into nanodiscs (CxI-NDs). a) Elution of CxI-NDs from the Superose 6 increase 5/150 size-exclusion column in the final step of the preparation (see Methods for details). The CxI-NDs elute at ∼1.6 mL, the same as the DDM-bound enzyme. b) Specific NADH:dQ activity data on CxI- DDM and CxI-NDs show that the complex I in CxI-NDs is highly catalytically competant, but catalysis is limited by substrate access. The reference activity (DDM-CxI, before reconstitution) is conserved in CxI-NDs following addition of CHAPS (to dissociate them) and asolectin (to create a lipidic phase). Without CHAPS/asolectin the activity was much lower, so either dQ does not exchange effectively in and out of the nanodisc, its mobility is restricted, or the MSPs prevent it from entering the active site. The low observed rate of NADH oxidation, which is further decreased in the absence of dQ, may either result from dQ reduction by complex I, or dQ-mediated reoxidation of the Q10H2 generated by complex I. Note that inhibitor-insensitive background rates have not been subtracted. 200 µM NADH was added in all cases (see Methods for experimental details).

**Supplementary Fig. 2:**
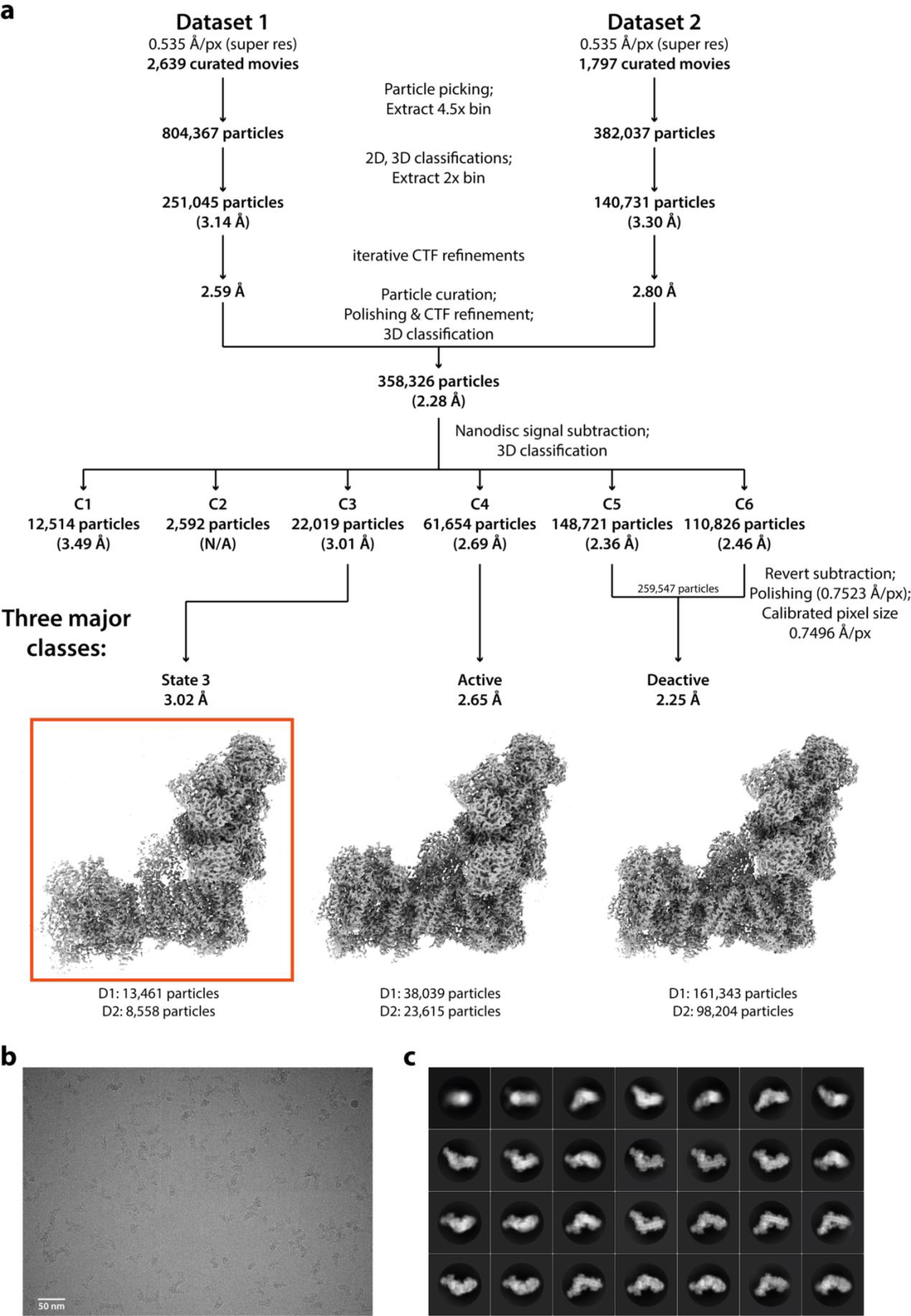
Cryo-EM data processing – global classification. a) A flow chart of cryo-EM data processing up to the final global 3D classification leading to the three major classes – active, deactive and state 3. D1 and D2 denote Datasets 1 and 2, respectively. Where applicable, the red box denotes that this is the final map for the class. b) A representative raw micrograph from the data collections. c) A representative 2D classification from the data processing.

**Supplementary Fig. 3:**
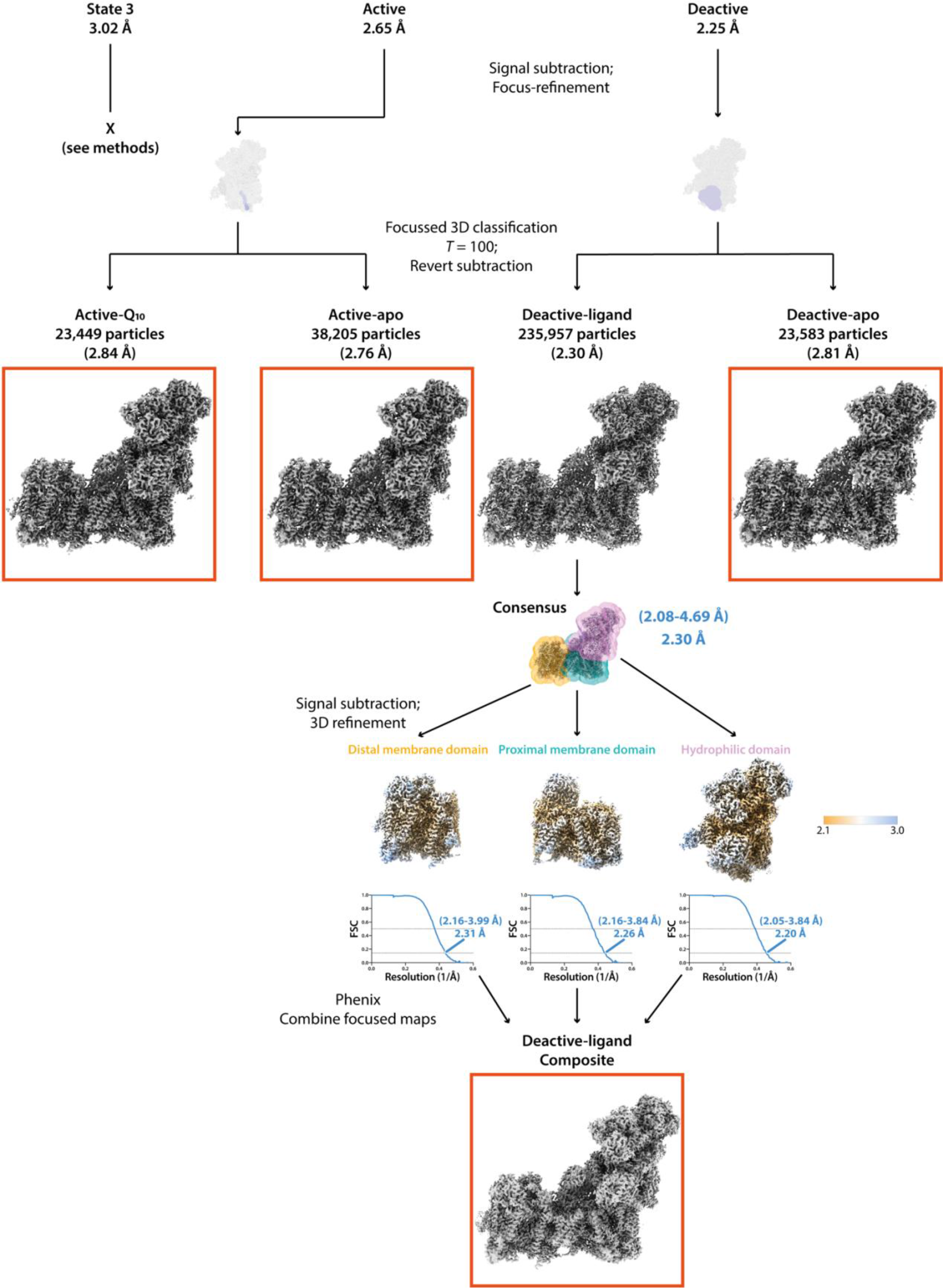
Cryo-EM data processing – local classification. A flow chart of cryo-EM data processing leading from the active and deactive major classes to their local substates (active- Q10, active-apo, deactive-ligand, and deactive-apo) by local classification. As substantial heterogeneity (apparent as weaker and/or discontinuous densities) was observed in the Q-binding sites of the active and deactive CxI-ND maps, the major classes were first signal subtracted to retain the hydrophilic arm, focus-refined, and then 3D focus-classified without alignment using class-specific local masks. A tight mask generated from a tentative Q10 model was used for local classification for the active class, and a more generous mask made from a provisional partial protein model (ND1, NDUFS2, and NDUFS7) encapsulating the Q-binding site region proved more successful for the deactive class. The consensus map for the deactive-ligand class showed poor densities at the distal region of the membrane arm due to movement in this region (subtle differences in the ‘openness’ of the hydrophilic and membrane arms and stronger alignment to the former mean that flexibility is exaggerated in the distal membrane domain). For this reason, a composite map was made for the deactive-ligand class. The consensus map was split into three regions by signal subtraction and focus-refinement using the three overlaid masks shown in transparent colours. Local resolution maps are shown for the distal (left) and proximal (middle) membrane domains, and the peripheral domain (right). Local resolutions were estimated using the Local Resolution function in RELION and plotted using UCSF ChimeraX^53^ with contour levels set to 0.015. The coloured key on the right indicates the resolution in Å, and the map resolution ranges are indicated in brackets for the respective maps, in blue. RELION half-map (sky blue) FSC curves are shown for each domain. The RELION map sharpening *B*-factors (Å^2^) for the consensus map, the distal and proximal membrane domains, and the peripheral domain were -58, -53, -49, and -51, respectively. The resulting composite map is shown in gray at contour level of 6.0. Composite maps were not made for the other classes either because there was little flexibility in the distal membrane domain (active-Q10 and active-apo) or because it did not lead to a marked improvement in map densities (deactive-apo and state 3). Where applicable, the red boxes denote the final maps for the classes.

**Supplementary Fig. 4:**
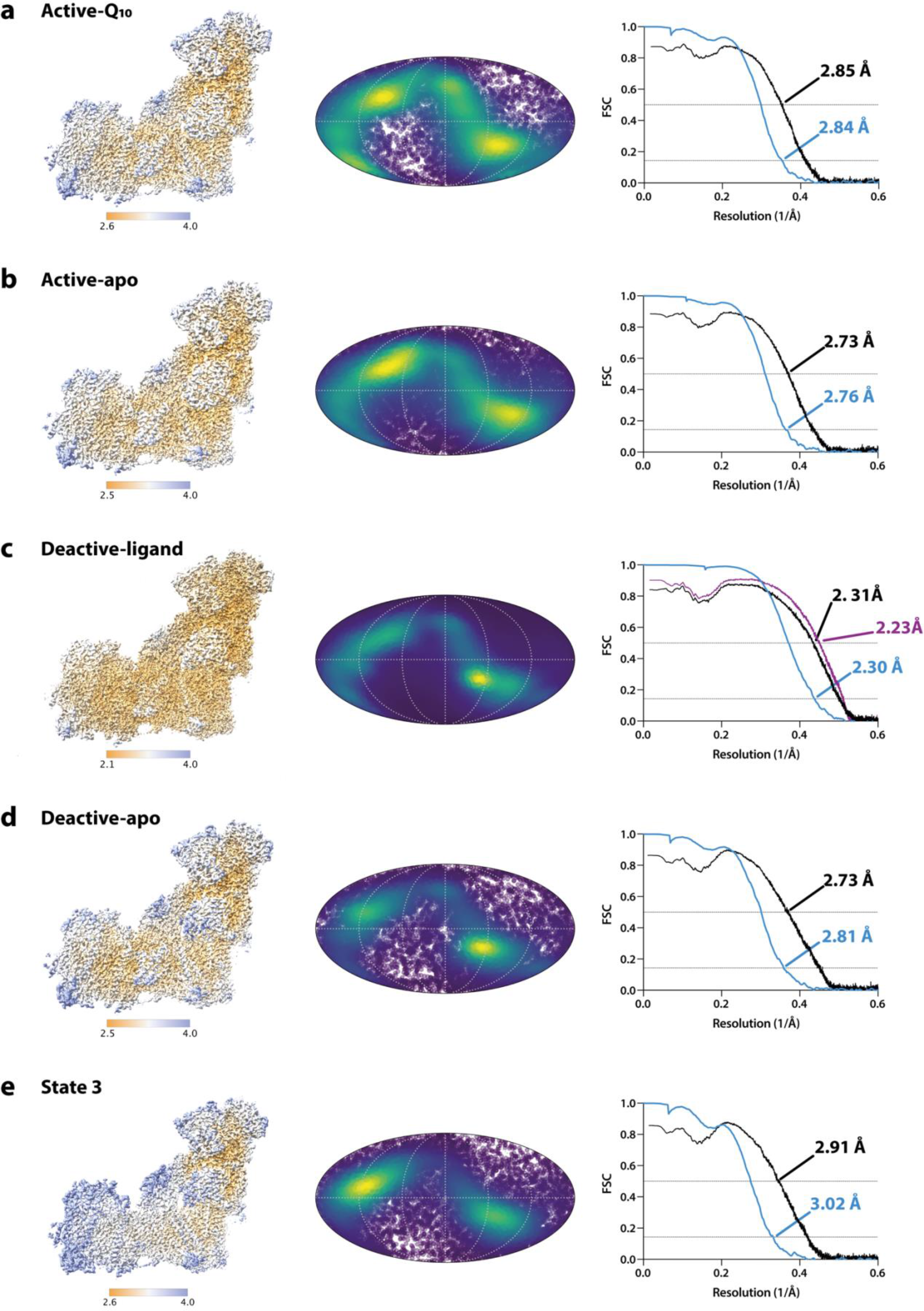
Local resolution maps, Mollweide projections, and Fourier shell correlation curves for the five CxI-ND states. Local resolution consensus maps (left), Mollweide projections (middle), and Fourier shell correlation (FSC) curves (right) are shown for the (a) active- Q10, (b) active-apo, (c) deactive-ligand, (d) deactive-apo, and (e) state 3 structures. The deactive- ligand map is the composite map described in Supplementary Fig. 3. Local resolutions were estimated using the Local Resolution function in RELION and plotted using UCSF ChimeraX with contour levels 6.5, 6.0, 4.0, 6.5, and 6.5, respectively. Coloured keys indicate resolution in Å. Mollweide projections were plotted using Python and Matplotlib. RELION half-map (sky blue) and model-map (black – locally-sharpened consensus maps; purple – focus-refined composite map) FSC curves are shown.

**Supplementary Fig. 5:**
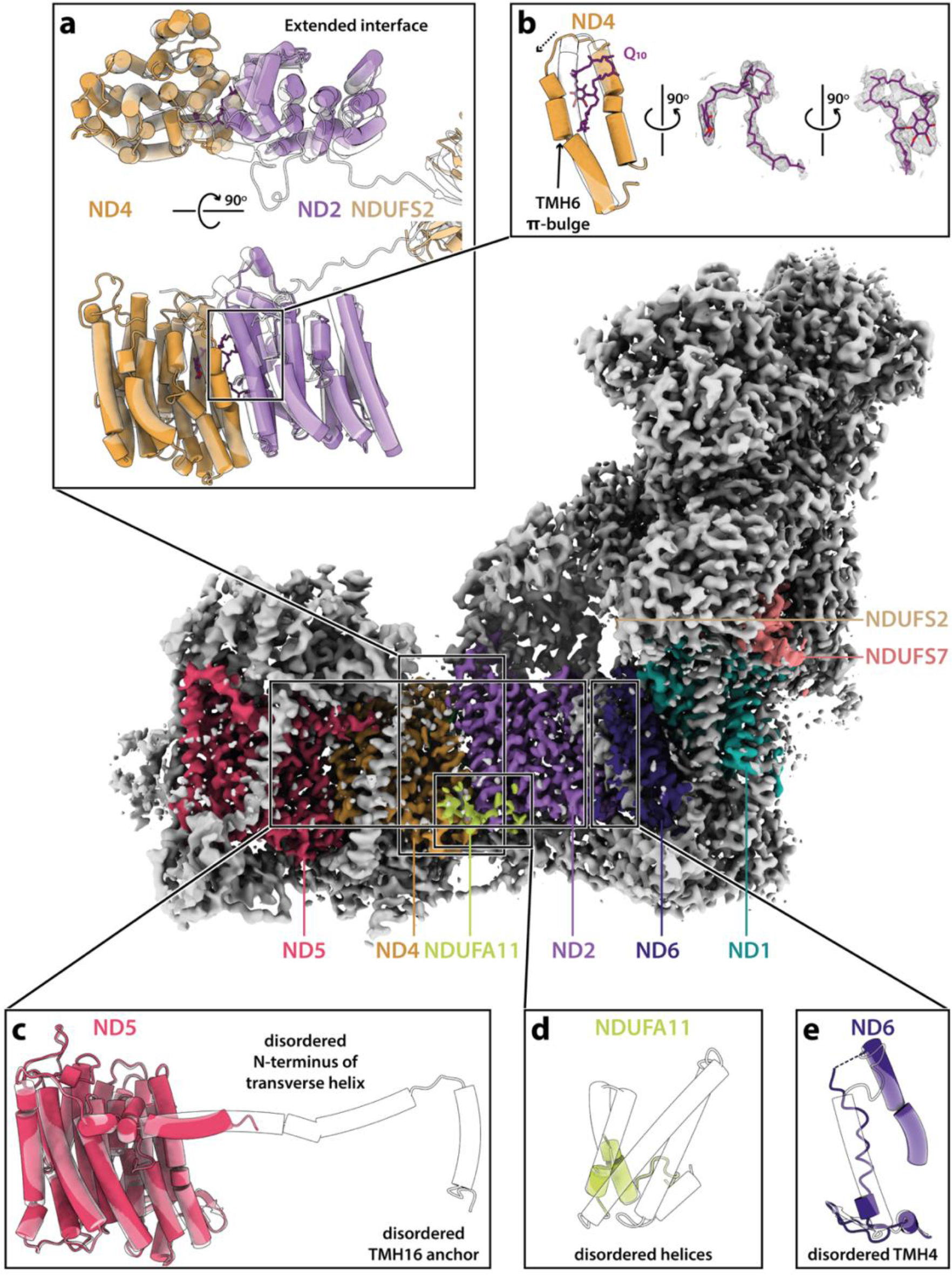
Structural features of state 3 of bovine complex I. The cryo-EM density map for state 3 is presented at a contour level of 6 in UCSF ChimeraX^53^. Cartoon and/or atomic representations are shown for (a) ND2, ND4, and NDUFS2, (b) ND4-TMH5-6 and Q10 (map contour level of 4.5), (c) ND5, (d) NDUFA11, and (e) ND6-TMH3-4. Models for state 3 are shown in solid colours and the deactive-ligand structure in transparent white with a black outline, aligned to the subunits shown. Conformational differences in the Q-binding site loops are shown in Fig. 4.

**Supplementary Fig. 6:**
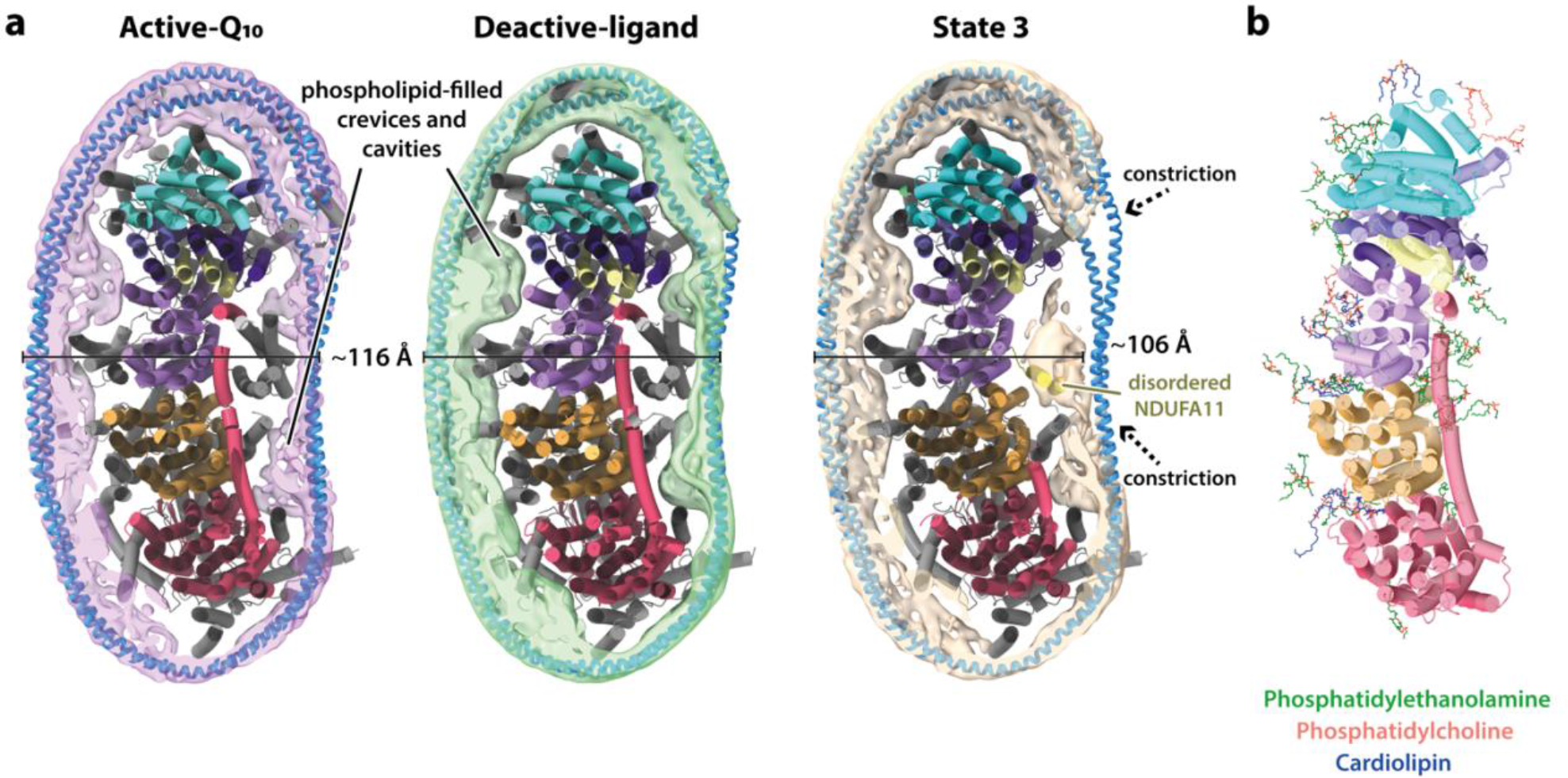
Structural features of complex I-reconstituted nanodiscs. a) Clipped top views of the active-Q10, deactive-ligand and state 3 CxI-ND structures (cartoon) and their subtract- refined MSP2N2 nanodiscs, with the MSP2N2 polyalanine model overlaid. Potential regions in the state 3 structure where the MSP2N2 helices may ‘wedge in’ are indicated with dashed arrows. Lateral nanodisc widths are indicated. b) 42 phospholipids modelled in a representative CxI-ND structure (deactive-ligand), with the seven core membrane subunits represented as cartoons. All non-cardiolipin phospholipids were modelled as phosphatidylethanolamines unless density features indicated phosphatidylcholine to be more likely. All subunits are coloured as in Fig. 1.

**Supplementary Fig. 7.**
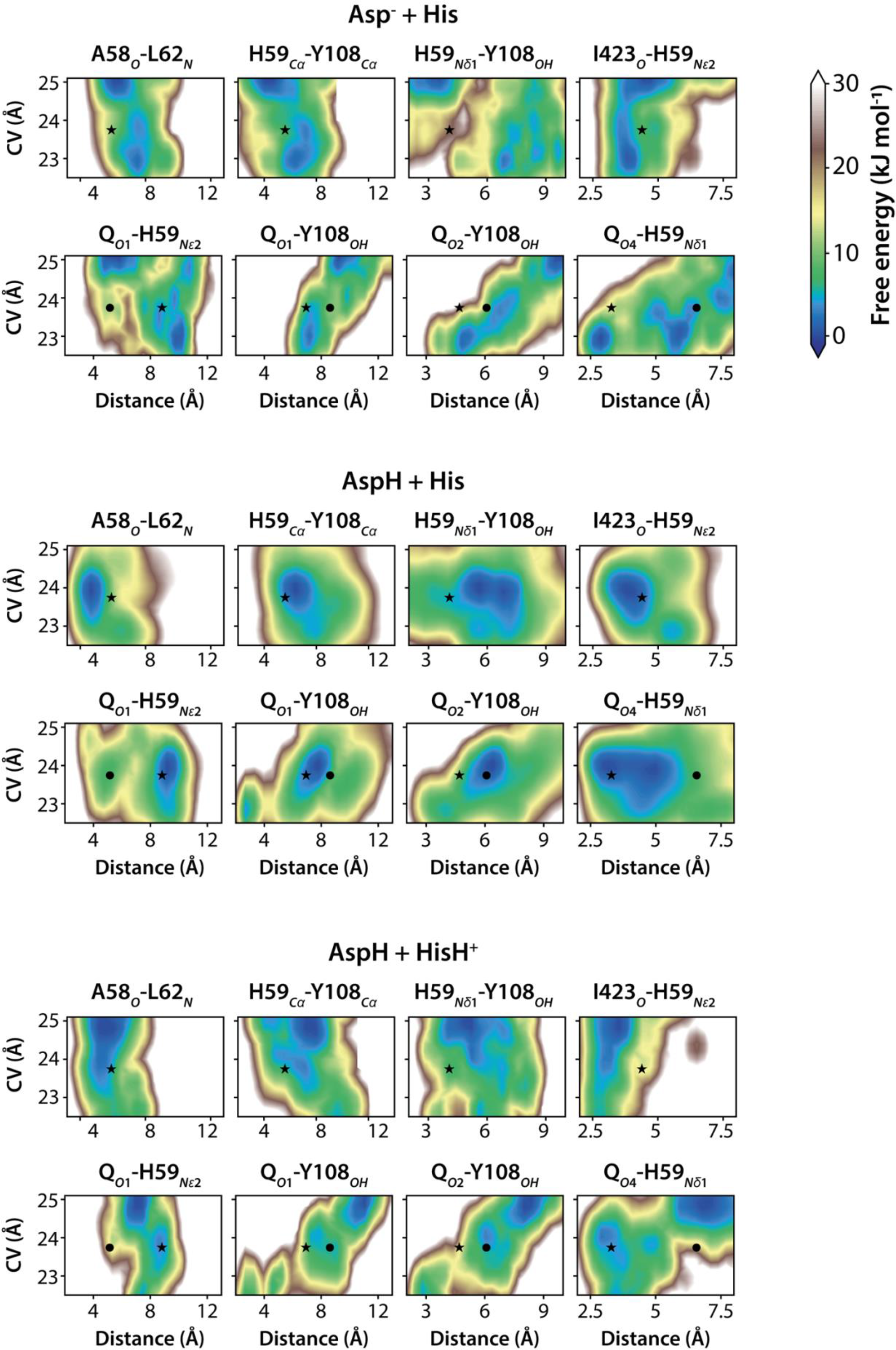
Free energy profiles for additional structural properties obtained from metadynamics simulations. Three combinations of sidechain protonation were simulated with Asp160^NDUFS2^ ionised (Asp^-^) or protonated (AspH) and His59^NDUFS2^ neutral (His, Nδ1-protonated π tautomer) or di-protonated (HisH^+^, both Nδ1- and Nε2-protonated), as indicated above each pair of panel rows. CV describes the Q-headgroup position along the binding channel (Fig 3b-d). Properties correspond to atom-pair distances shown on top of each column and symbols correspond to distances observed in active-Q10 cryo-EM models with primary (star) and flipped (bullet) Q- headgroup. The [AspH + His] charge-state displays free energy minima close (within 8 kJ mol^-1^) to distances observed in active-Q10 cryo-EM models for all structural properties. This is not true for the other two charge-states for distances involving protein centres, for example distance His59^NDUFS2^-Nδ1–Tyr108^NDUFS2^-OH for charge-state [Asp^-^ + His] and distance Ile423^NDUFS2^-O– His59^NDUFS2^-Nε2 for charge-state [AspH + HisH^+^], or involving the Q-headgroup, for example distances QO2–Tyr108^NDUFS2^-OH and QO4–His59^NDUFS2^-Nδ1 for charge-state [Asp^-^ + His], or distances QO1–Tyr108^NDUFS2^-OH and QO2–Tyr108^NDUFS2^-OH for charge-state [AspH + HisH^+^].

**Supplementary Fig. 8:**
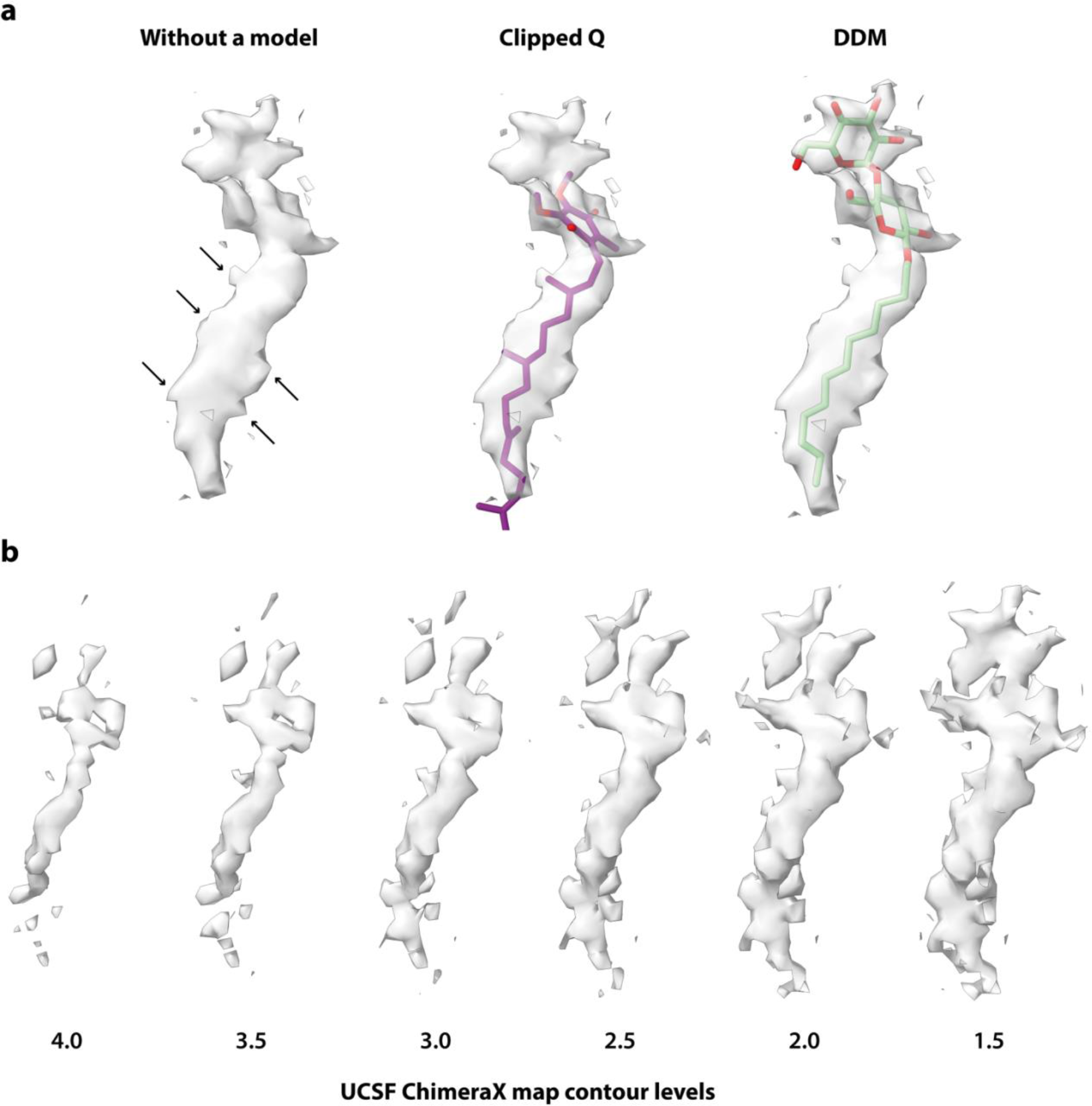
Features of the Coulomb potential density for the ligand in the deactive- ligand map. a) Focus-refined cryo-EM density of the ligand at contour level of 3 in an orthogonal view to Fig. 4b, showing zigzagged protrusions of the isoprenoid tail (indicated with arrows). Modelled Q (clipped) and DDM are shown. b) Locally-sharpened cryo-EM density of the ligand at decreasing map contour levels in UCSF ChimeraX^53^, showing the weaker density for the second headgroup gradually emerging.

**Supplementary Fig. 9:**
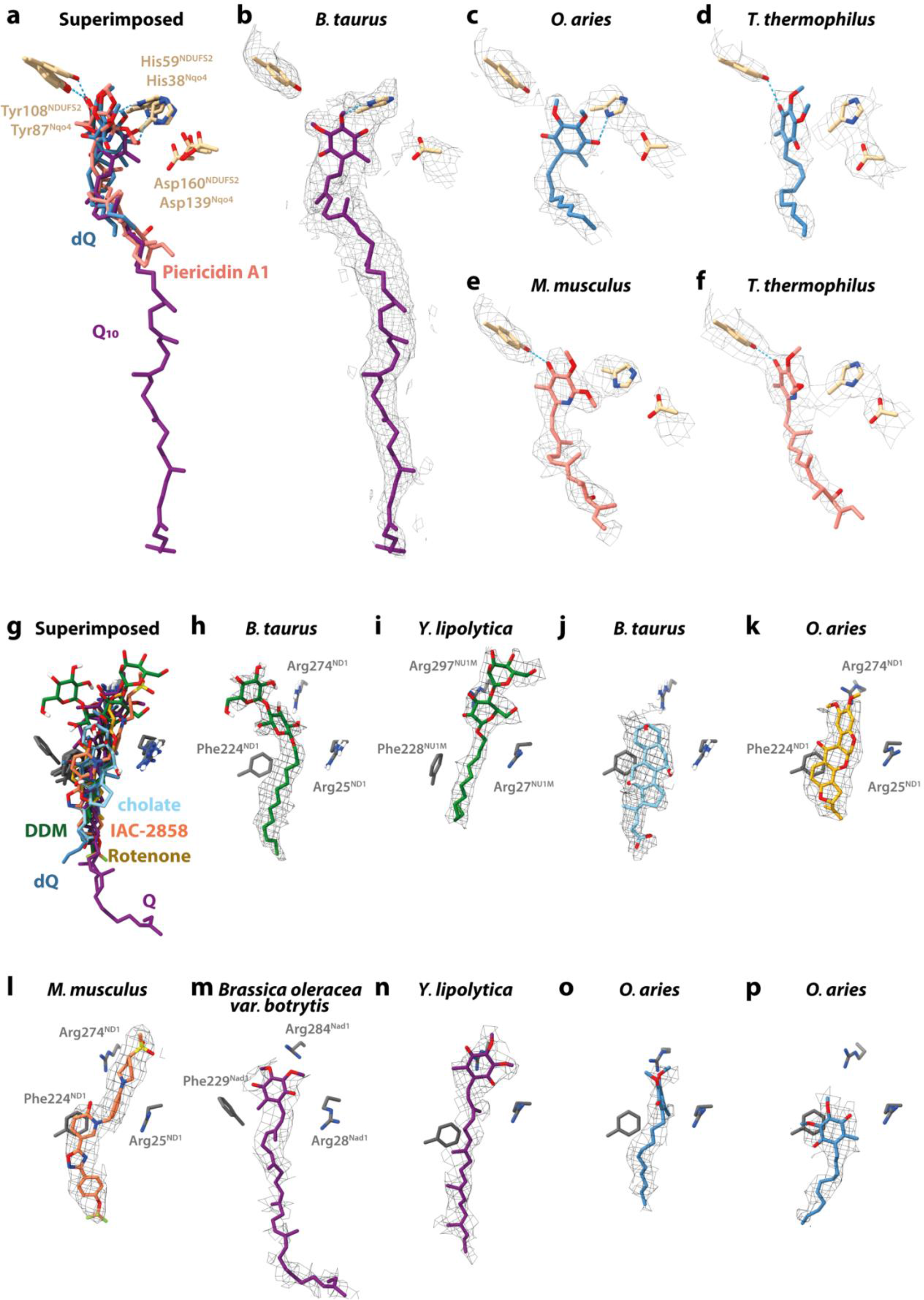
Comparisons between available cryo-EM and X-ray crystal structures of ligands bound in the Q-binding site of complex I. a-f) Q10, dQ, and piericidin A1 bound at the top of the Q-binding site. a) Superimposition of complex I-bound Q10, dQ and piericidin A1 molecules available from the PDB shown alongside Tyr108^NDUFS2^, His59^NDUFS2^, and Asp160^NDUFS2^ (mammalian numbering). b) Q10 bound in the bovine active-Q10 structure, with cryo-EM density shown at a contour level of 4.4 in UCSF ChimeraX^53^. c) dQ bound in ovine complex I in the ‘closed’ conformation reported to be undergoing turnover (PDB ID: 6ZKC), with cryo-EM density shown at contour level of 0.03 (EMD-11244)^22^. d) dQ bound in *T. thermophilus* complex I (PDB ID: 6I0D), with electron density shown at contour level of 0.2^21^. e) Piericidin A1 bound in mouse complex I (PDB ID: 6ZTQ), with cryo-EM density shown at contour level of 0.015 (EMD-11424)^12^. f) Piericidin A1 bound in *T. thermophilus* complex I (PDB ID: 6Q8O), with electron density shown at contour level of 0.06^21^. g-p) Various ligands bound at the entrance of the Q-binding site. g) Superimposition of complex I-bound DDM, cholate, rotenone, IACS-2858, Q9-10, and dQ molecules available from the PDB shown alongside Phe224^ND1^, Arg274^ND1^, and Arg25^ND1^ (mammalian numbering). h) DDM bound in the bovine deactive-ligand structure, with cryo-EM density shown at contour level of 4. i) DDM bound in *Y. lipolytica* complex I (PDB ID: 6YJ4), with cryo-EM density shown at contour level of 0.025 (EMD-10815)^25^. j) Cholate bound in the bovine state 3 structure, with cryo-EM density shown at contour level of 4.5. k) Rotenone bound in ovine complex I in the ‘open’ conformation (PDB ID: 6ZKM), with cryo-EM density shown at contour level of 0.05 (EMD-11254)^22^. l) IACS-2858 bound in murine complex I (PDB ID: 7B93), with cryo-EM density shown at contour level of 0.04 (EMD-12059)^23^. m) Q10 bound in *Brassica oleracea var. botrytis* complex I (PDB ID: 7A23), with cryo-EM density shown at contour level of 0.009 (EMD-11614)^29^. n) Q9 bound in *Y. lipolytica* complex I (PDB ID: 6RFR), with cryo-EM density shown at contour level of 0.01 (EMD-4873)^28^. o) dQ bound in ovine complex I in the ‘closed’ conformation reported to be undergoing turnover (PDB ID: 6ZKC), with cryo-EM density shown at contour level of 0.04 (EMD-11244)^22^. p) dQ bound in ovine complex I in the ‘open’ conformation reported to be undergoing turnover (PDB ID: 6ZKD), with cryo-EM density shown at contour level of 0.04 (EMD-11245)^22^.

## SUPPLEMENTARY INFORMATION

### Supplementary Discussion 1. Biochemical relevance of the state 3 structure

Improvements in resolution of the third state of complex I, state 3, now enable a more comprehensive consideration of this poorly understood class of particles (Supplementary Fig. 5). Previously, for state 3 we described a loss of clear density for the C-terminal half of the ND5 transverse helix and its anchor helix (TMH16), much of the adjacent NDUFA11 subunit, and the N-terminal loop of NDUFS2^5,11^. The resolution of our early data was low (5.6 Å), but the same characteristics are conserved in our current higher resolution map (3.0 Å), so it is clear that these structural elements are disordered and/or have dissociated from the structured binding locations that they occupy on the complex in the active and deactive states. Our data now reveals also that ND6-TMH4 has lost its α-helical secondary structure in state 3, instead appearing as a poorly ordered loop, on the same exposed side of the complex as NDUFA11 and ND5-TMH16, extending the region affected by the disorder. Comparison of the nanodisc structures also shows clear differences in this region (Supplementary Fig. 6). Instead of clear densities stretching around NDUFA11 (as observed in the active and deactive states), the MSP2N2 helices in state 3 are not clearly resolved, consistent with substantial disorder in subunit NDUFA11 and/or with the MSP2N2 helices contracting inwards to occupy the space where NDUFA11 is usually observed. This change in nanodisc structure suggests that state 3 was present in the preparation before the enzyme was reconstituted into the nanodiscs. Therefore, it may result from destabilisation of the detergent-solubilised enzyme during its purification, but is not an artefact from cryo-EM grid preparation due to instability induced at the air-water interface^85–87^. Interestingly, NDUFA11 has been reported to be the only subunit in the membrane arm of complex I that undergoes rapid protein turnover in C2C12 myotubes^88^, consistent with a higher propensity to dissociate *in vivo*. Furthermore, in the mammalian respirasome NDUFA11 is sandwiched in-between complex III and the core subunits of complex I^89–91^, and the removal of complex III during solubilisation of the membrane may introduce additional instability to NDUFA11 and adjacent structures.

Comparing the three states of complex I resolved here shows clearly that state 3 has more in common with the deactive state than the active state. Globally, the apparent angle between the hydrophilic and hydrophobic domains of the complex is most acute in the active state, considerably more obtuse in the deactive state and similar, or greater still, in state 3. Key structural features that distinguish the deactive state from the active state are also present in state 3, including the π-bulge in ND6-TMH3, disordered ND3 TMH1-2 loop and structures around NDUFS7-Arg77. In contrast, the NDUFS2-β1-β2 and ND1 TMH5-6 loop conformations differ from those observed in both the active and deactive states. However, as features of the Q-binding site are known to be remodelled by ligands, it is unclear whether their conformations are a feature of state 3 or determined by ligand binding. Finally, the features discussed above that are disordered in state 3 are clearly ordered (and equivalent) in both the active and deactive states, and the ND2-ND4 interface is closed in both those states. Therefore, state 3 shares a substantial number of distinguishing features with the deactive state – but it shares none of them with the active state. Due to the extra-relaxed and disordered characteristics of state 3, we propose to name this state ‘slack’ complex I in the future.

The three ‘open’ states described for native ovine complex I^22^ exhibit all the key structural features observed in our deactive states, but also features that are specific to our state 3 structure. In particular, the state named open3 displays extensive disorder (evident as weak or absent densities) for NDUFA11, the C-terminal half of the ND5 transverse helix and its anchor helix (TMH16), the ∼40 N-terminal residues of NDUFS2, and ND6-TMH4. These comparisons suggest that the open conformations of ovine complex I exist on a structural spectrum between our deactive state and state 3. Similar state 3-like characteristics are also observed in maps for ovine complex I reported to be in the deactive state, which we note was prepared by incubating a concentrated aliquot of the detergent-solubilised complex I at 32 °C for 30 min after anion exchange^22^, rather than by deactivating the enzyme in its stabilising native membrane environment^9, 10, 13^.

In state 3 the extended interface between subunits ND2 and ND4 contains an ordered Q10 molecule, accommodated by a π-bulge formed in ND4-TMH6. It is possible that the ND4-TMH6 π-bulge forms during catalysis, such that state 3 represents a relaxed state of a hitherto-uncharacterised intermediate, in which the π-bulge causes Tyr148^ND4^ to pivot away from its usual hydrogen bond to Glu559^ND5^. By doing so it may destabilise the ND2-ND4 interface, propagating instability to the transverse helix and associated structures, resulting in the observed loss of structural integrity. The interfaces between NDUFA10, ND4, and ND2 have also been observed to accumulate strain in simulation work, and proposed to modulate conformational changes within the membrane domain^46^. Intriguingly, the position of the Q10-headgroup bound at the ND2-ND4 interface in state 3 coincides with the position of a rotenone inhibitor molecule detected previously in ovine complex I open conformations^22^. Although it is difficult to conceive of any functional role for Q10 binding at this site, so distant from any of the enzyme’s redox cofactors, it is possible that any molecule binding here could lock the position of Tyr148^ND4^ and thereby be inhibitory – if reorganisation of ND4-TMH6 really does occur on the catalytic cycle.

Loosening the structural constraints of the transverse helix in state 3 and opening up of the ND2- ND4 interface may alternatively allow the Q10/rotenone molecule to enter in adventitiously. The propensity of the enzyme to fracture at this interface is highlighted by treatment of bovine complex I with the zwitterionic detergent *N,N*-dimethyldodecylamine *N*-oxide (LDAO) to form the ND4-ND5- containing subcomplex of the distal membrane domain known as subcomplex Iβ^92^. The same fragmentation occurred during crystallisation trials on the DDM-solubilised enzyme^93^, yielding a structure of subcomplex Iβ^92^ in which the C-terminal section of ND5 is highly disordered. The tendency of the ND4-ND5 module to dissociate was also illustrated by *Y. lipolytica* knock-out strain *nb8m*Δ (where NB8M is a homologue of subunit NDUFB7, adjacent to the C-terminus of ND5), in which the distal membrane domain was absent^94^. These observations argue that state 3 may represent an enzyme in the initial stages of degradation, not a catalytically relevant state.

To define the catalytic capability of state 3 it will be necessary to produce a homogeneous sample of it and characterise it both functionally and structurally. It is currently unclear how to achieve this for bovine complex I, but we note that a structure we solved previously for complex I from rhesus macaque (*Macaca mulatta*)^11^ was dominated by particles in this state (lacking clear densities for NDUFA11, the C-terminal half of the ND5 transverse helix and its anchor helix (TMH16), the N-terminus of NDUFS2, and parts of ND6-TMH4) – and the preparation exhibited only a very low activity of 0.5 µmol min^-1^ mg^-1^ ^11^. Current evidence to suggest that state 3-like states do exist on the catalytic cycle is limited to their presence in a sample of ovine complex I treated with the substrates for turnover – however, similar states were present in the starting mixture and may simply not have responded to the addition of substrates^22^. Further investigations of complex I samples under established turnover conditions and in defined biochemical states will be necessary to finally establish the relevance of this intriguing enzyme state.

### Supplementary Discussion 2: Minor differences between the structures of nanodisc-reconstituted and DDM-solubilised complex I

First, in all CxI-ND maps, the tip of the β-hairpin in the ND6 TMH4-5 loop on the intermembrane space face is shifted by 3-3.5 Å towards subunit ND2, so that it makes contact with the ND2-TMH4- 5 helix-loop-helix motif and ND4L-TMH1. In the DDM-solubilised reference active-state structure^43^, there is an unmodelled density between the two structural elements that resembles a DDM molecule displacing the ND6-β-hairpin. The β-hairpin motif is not present in *M. musculus* complex I^10, 13^, while in the lauryl maltose neopentyl glycol (LMNG)-solubilised ovine enzyme^22^ its conformation matches the CxI-ND conformation.

Second, a density for a steroid molecule (modelled as a cholate, which was added at the reconstitution step) is present in all CxI-ND maps at the interface between subunits ND5 and NDUFB6. The nearby NDUFB6 matrix helix/loop is resolved for the first time here, perhaps as a result of the structure being stabilised by incorporation of the additional cholate molecule and enclosed by the adjacent MSP2N2 helices – although the previously unstructured region makes remarkably few direct contacts with any other subunits.

Third, the C-terminal peptide of mammalian subunit NDUFS1 (ca. 10 residues) is also resolved for the first time here, with the terminal Cys704^NDUFS1^ residue lodged between subunits NDUFS1, NDUFS6, and NDUFS8. The absence of DDM in the sample buffer may be responsible for the improved resolution of this peptide in the nanodisc structures, on the outside of the hydrophilic domain.

## Notes

### Competing Interest Statement

The authors have declared no competing interest.

## References

1. Hirst, J. Mitochondrial Complex I. Annu. Rev. Biochem. 82, 551–575 (2013).

2. Parey, K., Wirth, C., Vonck, J. & Zickermann, V. Respiratory complex I — structure, mechanism and evolution. Curr. Opin. Struct. Biol. 63, 1–9 (2020).

3. Hirst, J., Carroll, J., Fearnley, I. M., Shannon, R. J. & Walker, J. E. The nuclear encoded subunits of complex I from bovine heart mitochondria. Biochim. Biophys. Acta - Bioenerg. 1604, 135–150 (2003).

4. Stroud, D. A. et al. Accessory subunits are integral for assembly and function of human mitochondrial complex I. Nature 538, 123–126 (2016).

5. Zhu, J., Vinothkumar, K. R. & Hirst, J. Structure of mammalian respiratory complex I. Nature 536, 354–358 (2016).

6. Murphy, M. P. & Hartley, R. C. Mitochondria as a therapeutic target for common pathologies. Nat. Rev. Drug Discov. 17, 865–886 (2018).

7. Fassone, E. & Rahman, S. Complex I deficiency: clinical features, biochemistry and molecular genetics. J. Med. Genet. 49, 578–590 (2012).

8. Fiedorczuk, K. & Sazanov, L. A. Mammalian Mitochondrial Complex I Structure and Disease-Causing Mutations. Trends Cell Biol. 28, 835–867 (2018).

9. Blaza, J. N., Vinothkumar, K. R. & Hirst, J. Structure of the Deactive State of Mammalian Respiratory Complex I. Structure 26, 312–319 (2018).

10. Agip, A.-N. A. et al. Cryo-EM structures of complex I from mouse heart mitochondria in two biochemically defined states. Nat. Struct. Mol. Biol. 25, 548–556 (2018).

11. Agip, A.-N. A., Blaza, J. N., Fedor, J. G. & Hirst, J. Mammalian Respiratory Complex I Through the Lens of Cryo-EM. Annu. Rev. Biophys. 48, 165–184 (2019).

12. Bridges, H. R. et al. Structure of inhibitor-bound mammalian complex I. Nat. Commun. 11, 5261 (2020).

13. Yin, Z. et al. Structural basis for a complex I mutation that blocks pathological ROS production. Nat. Commun. 12, 707 (2021).

14. Kotlyar, A. B. & Vinogradov, A. D. Slow active/inactive transition of the mitochondrial NADH-ubiquinone reductase. Biochim. Biophys. Acta - Bioenerg. 1019, 151–158 (1990).

15. Vinogradov, A. D. Catalytic properties of the mitochondrial NADH–ubiquinone oxidoreductase (Complex I) and the pseudo-reversible active/inactive enzyme transition. Biochim. Biophys. Acta - Bioenerg. 1364, 169–185 (1998).

16. Babot, M., Birch, A., Labarbuta, P. & Galkin, A. Characterisation of the active/de-active transition of mitochondrial complex I. Biochim. Biophys. Acta - Bioenerg. 1837, 1083–1092 (2014).

17. Galkin, A. & Moncada, S. Modulation of the conformational state of mitochondrial complex I as a target for therapeutic intervention. Interface Focus 7, 20160104 (2017).

18. Galkin, A., Abramov, A. Y., Frakich, N., Duchen, M. R. & Moncada, S. Lack of Oxygen Deactivates Mitochondrial Complex I. J. Biol. Chem. 284, 36055–36061 (2009).

19. Chouchani, E. T. et al. Cardioprotection by S-nitrosation of a cysteine switch on mitochondrial complex I. Nat. Med. 19, 753–759 (2013).

20. Zickermann, V. et al. Mechanistic insight from the crystal structure of mitochondrial complex I. Science 347, 44–49 (2015).

21. Gutiérrez-Fernández, J. et al. Key role of quinone in the mechanism of respiratory complex I. Nat. Commun. 11, 4135 (2020).

22. Kampjut, D. & Sazanov, L. A. The coupling mechanism of mammalian respiratory complex I. Science 370, eabc4209 (2020).

23. Chung, I. et al. Cork-in-bottle mechanism of inhibitor binding to mammalian complex I. Sci. Adv. 7, eabg4000 (2021).

24. Grba, D. N. et al. Cryo-electron microscopy reveals how acetogenins inhibit mitochondrial respiratory complex I. J. Biol. Chem. (2022) doi:10.1016/j.jbc.2022.101602.

25. Grba, D. N. & Hirst, J. Mitochondrial complex I structure reveals ordered water molecules for catalysis and proton translocation. Nat. Struct. Mol. Biol. 27, 892–900 (2020).

26. Parey, K. et al. Cryo-EM structure of respiratory complex I at work. Elife 7, 1–28 (2018).

27. Parey, K. et al. High-resolution structure and dynamics of mitochondrial complex I— Insights into the proton pumping mechanism. Sci. Adv. 7, eabj3221 (2021).

28. Parey, K. et al. High-resolution cryo-EM structures of respiratory complex I: Mechanism, assembly, and disease. Sci. Adv. 5, eaax9484 (2019).

29. Soufari, H., Parrot, C., Kuhn, L., Waltz, F. & Hashem, Y. Specific features and assembly of the plant mitochondrial complex I revealed by cryo-EM. Nat. Commun. 11, 5195 (2020).

30. Klusch, N., Senkler, J., Yildiz, Ö., Kühlbrandt, W. & Braun, H.-P. A ferredoxin bridge connects the two arms of plant mitochondrial complex I. Plant Cell 33, 2072–2091 (2021).

31. Baradaran, R., Berrisford, J. M., Minhas, G. S. & Sazanov, L. A. Crystal structure of the entire respiratory complex I. Nature 494, 443–448 (2013).

32. Tocilescu, M. A. et al. The role of a conserved tyrosine in the 49-kDa subunit of complex I for ubiquinone binding and reduction. Biochim. Biophys. Acta - Bioenerg. 1797, 625–632 (2010).

33. Gamiz-Hernandez, A. P., Jussupow, A., Johansson, M. P. & Kaila, V. R. I. Terminal Electron– Proton Transfer Dynamics in the Quinone Reduction of Respiratory Complex I. J. Am. Chem. Soc. 139, 16282–16288 (2017).

34. Warnau, J. et al. Redox-coupled quinone dynamics in the respiratory complex I. Proc. Natl. Acad. Sci. 115, E8413–E8420 (2018).

35. Teixeira, M. H. & Arantes, G. M. Balanced internal hydration discriminates substrate binding to respiratory complex I. Biochim. Biophys. Acta - Bioenerg. 1860, 541–548 (2019).

36. Röpke, M. et al. Deactivation blocks proton pathways in the mitochondrial complex I. Proc. Natl. Acad. Sci. 118, e2019498118 (2021).

37. Fedor, J. G., Jones, A. J. Y., Di Luca, A., Kaila, V. R. I. & Hirst, J. Correlating kinetic and structural data on ubiquinone binding and reduction by respiratory complex I. Proc. Natl. Acad. Sci. 114, 12737–12742 (2017).

38. Ritchie, T. K. et al. Reconstitution of Membrane Proteins in Phospholipid Bilayer Nanodiscs. in Methods in Enzymology vol. 464 211–231 (Academic Press, 2009).

39. Biner, O., Fedor, J. G., Yin, Z. & Hirst, J. Bottom-Up Construction of a Minimal System for Cellular Respiration and Energy Regeneration. ACS Synth. Biol. 9, 1450–1459 (2020).

40. Zivanov, J., Nakane, T. & Scheres, S. H. W. Estimation of high-order aberrations and anisotropic magnification from cryo-EM data sets in RELION -3.1. IUCrJ 7, 253–267 (2020).

41. Scheres, S. H. W. Processing of Structurally Heterogeneous Cryo-EM Data in RELION. In Methods in Enzymology vol. 579 125–157 (Elsevier Inc., 2016).

42. Bibow, S. et al. Solution structure of discoidal high-density lipoprotein particles with a shortened apolipoprotein A-I. Nat. Struct. Mol. Biol. 24, 187–193 (2017).

43. Bridges, H. R., Blaza, J. N., Yin, Z., Chung, I. & Hirst, J. Bovine complex I in the active state at 3.1 Å. (2022) doi:10.2210/pdb7QSD/pdb.

44. Marques, M. A., Purdy, M. D. & Yeager, M. CryoEM maps are full of potential. Curr. Opin. Struct. Biol. 58, 214–223 (2019).

45. Burger, N. et al. ND3 Cys39 in complex I is exposed during mitochondrial respiration. Cell Chem. Biol. (2021) doi:10.1016/j.chembiol.2021.10.010.

46. Di Luca, A. & Kaila, V. R. I. Molecular strain in the active/deactive-transition modulates domain coupling in respiratory complex I. Biochim. Biophys. Acta - Bioenerg. 1862, 148382 (2021).

47. Tian, W., Chen, C., Lei, X., Zhao, J. & Liang, J. CASTp 3.0: computed atlas of surface topography of proteins. Nucleic Acids Res. 46, W363–W367 (2018).

48. Taguchi, A. T., Mattis, A. J., O’Malley, P. J., Dikanov, S. A. & Wraight, C. A. Tuning Cofactor Redox Potentials: The 2-Methoxy Dihedral Angle Generates a Redox Potential Difference of >160 mV between the Primary (QA) and Secondary (QB) Quinones of the Bacterial Photosynthetic Reaction Center. Biochemistry 52, 7164–7166 (2013).

49. Haapanen, O., Djurabekova, A. & Sharma, V. Role of second quinone binding site in proton pumping by respiratory complex I. Front. Chem. 7, 1–14 (2019).

50. Wang, P., Dhananjayan, N., Hagras, M. A. & Stuchebrukhov, A. A. Respiratory complex I: Bottleneck at the entrance of quinone site requires conformational change for its opening. Biochim. Biophys. Acta - Bioenerg. 1862, 148326 (2021).

51. Dhananjayan, N., Wang, P., Leontyev, I. & Stuchebrukhov, A. A. Quinone binding in respiratory complex I: Going through the eye of a needle. The squeeze-in mechanism of passing the narrow entrance of the quinone site. Photochem. Photobiol. Sci. 21, 1–12 (2022).

52. Sharma, V. et al. Redox-induced activation of the proton pump in the respiratory complex I. Proc. Natl. Acad. Sci. 112, 11571–11576 (2015).

53. Goddard, T. D. et al. UCSF ChimeraX: Meeting modern challenges in visualization and analysis. Protein Sci. 27, 14–25 (2018).

## Methods References

54. Blaza, J. N., Serreli, R., Jones, A. J. Y., Mohammed, K. & Hirst, J. Kinetic evidence against partitioning of the ubiquinone pool and the catalytic relevance of respiratory-chain supercomplexes. Proc. Natl. Acad. Sci. 111, 15735–15740 (2014).

55. Nagle, J. F. & Tristram-Nagle, S. Structure of lipid bilayers. Biochim. Biophys. Acta - Rev. Biomembr. 1469, 159–195 (2000).

56. Russo, C. J. & Passmore, L. A. Ultrastable gold substrates for electron cryomicroscopy. Science 346, 1377–1380 (2014).

57. Rohou, A. & Grigorieff, N. CTFFIND4: Fast and accurate defocus estimation from electron micrographs. J. Struct. Biol. 192, 216–221 (2015).

58. Zivanov, J. et al. New tools for automated high-resolution cryo-EM structure determination in RELION-3. Elife 7, e42166 (2018).

59. Pettersen, E. F. et al. UCSF Chimera—A visualization system for exploratory research and analysis. J. Comput. Chem. 25, 1605–1612 (2004).

60. Liebschner, D. et al. Macromolecular structure determination using X-rays, neutrons and electrons: recent developments in Phenix. Acta Crystallogr. Sect. D Struct. Biol. 75, 861– 877 (2019).

61. Emsley, P. & Cowtan, K. Coot: model-building tools for molecular graphics. Acta Crystallogr. Sect. D Biol. Crystallogr. 60, 2126–2132 (2004).

62. Fearnley, I. M., Finel, M., Skehel, J. M. & Walker, J. E. NADH:ubiquinone oxidoreductase from bovine heart mitochondria. cDNA sequences of the import precursors of the nuclear- encoded 39 kDa and 42 kDa subunits. Biochem. J. 278, 821–829 (1991).

63. Carroll, J., Fearnley, I. M., Shannon, R. J., Hirst, J. & Walker, J. E. Analysis of the Subunit Composition of Complex I from Bovine Heart Mitochondria. Mol. Cell. Proteomics 2, 117– 126 (2003).

64. Pintilie, G. et al. Measurement of atom resolvability in cryo-EM maps with Q-scores. Nat. Methods 17, 328–334 (2020).

65. Croll, T. I. ISOLDE: a physically realistic environment for model building into low-resolution electron-density maps. Acta Crystallogr. Sect. D Struct. Biol. 74, 519–530 (2018).

66. Scheres, S. H. W. RELION: Implementation of a Bayesian approach to cryo-EM structure determination. J. Struct. Biol. 180, 519–530 (2012).

67. Schrodinger LLC. The PyMOL Molecular Graphics System, Version 2.4.1. (2021).

68. Mills, J. E. J. & Dean, P. M. Three-dimensional hydrogen-bond geometry and probability information from a crystal survey. J. Comput. Aided. Mol. Des. 10, 607–622 (1996).

69. Meng, E. C. & Lewis, R. A. Determination of molecular topology and atomic hybridization states from heavy atom coordinates. J. Comput. Chem. 12, 891–898 (1991).

70. Olsson, M. H. M., Søndergaard, C. R., Rostkowski, M. & Jensen, J. H. PROPKA3: Consistent Treatment of Internal and Surface Residues in Empirical p*K*a Predictions. J. Chem. Theory Comput. 7, 525–537 (2011).

71. Abraham, M. J. et al. GROMACS: High performance molecular simulations through multi- level parallelism from laptops to supercomputers. SoftwareX 1–2, 19–25 (2015).

72. Javanainen, M. Universal Method for Embedding Proteins into Complex Lipid Bilayers for Molecular Dynamics Simulations. J. Chem. Theory Comput. 10, 2577–2582 (2014).

73. Teixeira, M. H. & Arantes, G. M. Effects of lipid composition on membrane distribution and permeability of natural quinones. RSC Adv. 9, 16892–16899 (2019).

74. Branduardi, D., Gervasio, F. L. & Parrinello, M. From A to B in free energy space. J. Chem. Phys. 126, 054103 (2007).

75. Barducci, A., Bussi, G. & Parrinello, M. Well-Tempered Metadynamics: A Smoothly Converging and Tunable Free-Energy Method. Phys. Rev. Lett. 100, 020603 (2008).

76. Klauda, J. B. et al. Update of the CHARMM All-Atom Additive Force Field for Lipids: Validation on Six Lipid Types. J. Phys. Chem. B 114, 7830–7843 (2010).

77. Huang, J. & MacKerell, A. D. CHARMM36 all-atom additive protein force field: Validation based on comparison to NMR data. J. Comput. Chem. 34, 2135–2145 (2013).

78. Huang, J. et al. CHARMM36m: an improved force field for folded and intrinsically disordered proteins. Nat. Methods 14, 71–73 (2017).

79. Jorgensen, W. L., Chandrasekhar, J., Madura, J. D., Impey, R. W. & Klein, M. L. Comparison of simple potential functions for simulating liquid water. J. Chem. Phys. 79, 926–935 (1983).

80. Chang, C. H. & Kim, K. Density Functional Theory Calculation of Bonding and Charge Parameters for Molecular Dynamics Studies on [FeFe] Hydrogenases. J. Chem. Theory Comput. 5, 1137–1145 (2009).

81. McCullagh, M. & Voth, G. A. Unraveling the Role of the Protein Environment for [FeFe]- Hydrogenase: A New Application of Coarse-Graining. J. Phys. Chem. B 117, 4062–4071 (2013).

82. Galassi, V. V. & Arantes, G. M. Partition, orientation and mobility of ubiquinones in a lipid bilayer. Biochim. Biophys. Acta - Bioenerg. 1847, 1560–1573 (2015).

83. Darden, T., York, D. & Pedersen, L. Particle mesh Ewald: An *N*⋅log(*N*) method for Ewald sums in large systems. J. Chem. Phys. 98, 10089–10092 (1993).

84. Tribello, G. A., Bonomi, M., Branduardi, D., Camilloni, C. & Bussi, G. PLUMED 2: New feathers for an old bird. Comput. Phys. Commun. 185, 604–613 (2014).

## Supplementary References

85. Glaeser, R. M. & Han, B.-G. Opinion: hazards faced by macromolecules when confined to thin aqueous films. Biophys. Reports 3, 1–7 (2017).

86. Noble, A. J. et al. Routine single particle CryoEM sample and grid characterization by tomography. Elife 7, 1–42 (2018).

87. D’Imprima, E. et al. Protein denaturation at the air-water interface and how to prevent it. Elife 8, 1–18 (2019).

88. Krishna, S. et al. Identification of long-lived proteins in the mitochondria reveals increased stability of the electron transport chain. Dev. Cell 56, 2952–2965 (2021).

89. Gu, J. et al. The architecture of the mammalian respirasome. Nature 537, 639–643 (2016).

90. Wu, M., Gu, J., Guo, R., Huang, Y. & Yang, M. Structure of Mammalian Respiratory Supercomplex I1III2IV1. Cell 167, 1598–1609 (2016).

91. Letts, J. A., Fiedorczuk, K., Degliesposti, G., Skehel, M. & Sazanov, L. A. Structures of Respiratory Supercomplex I+III2 Reveal Functional and Conformational Crosstalk. Mol. Cell 75, 1131–1146 (2019).

92. Sazanov, L. A., Peak-Chew, S. Y., Fearnley, I. M. & Walker, J. E. Resolution of the Membrane Domain of Bovine Complex I into Subcomplexes: Implications for the Structural Organization of the Enzyme. Biochemistry 39, 7229–7235 (2000).

93. Zhu, J. et al. Structure of subcomplex Iβ of mammalian respiratory complex I leads to new supernumerary subunit assignments. Proc. Natl. Acad. Sci. 112, 12087–12092 (2015).

94. Dröse, S. et al. Functional Dissection of the Proton Pumping Modules of Mitochondrial Complex I. PLoS Biol. 9, e1001128 (2011).

